# Retrograde Transport and ATG-4.2-Mediated Maturation Cooperate to Remove Autophagosomes from the Synapse

**DOI:** 10.1101/287144

**Authors:** Sarah E. Hill, Daniel A. Colón-Ramos

## Abstract

Autophagy is spatially compartmentalized in neurons, with autophagosome biogenesis occurring in the axon and degradation in the cell body. The mechanisms that coordinate autophagosome formation, trafficking and degradation across the polarized structure of the neuron are not well understood. Here we use genetic screens and *in vivo* imaging in single neurons of *C. elegans* to demonstrate that specific steps of autophagy are differentially required in distinct subcellular compartments of the neuron. We demonstrate that completion of autophagosome biogenesis and closure at the synapse are necessary for dynein-mediated retrograde transport. We uncover a role for UNC-16/JIP3/Sunday Driver in facilitating autophagosome retrograde transport. Through forward genetic screens we then determine that autophagosome maturation and degradation in the cell body depend on removal of LGG-1/Atg8/GABARAP from autophagosomes by the protease ATG-4.2. Our studies reveal that regulation of distinct ATG4 proteases contributes to the coordination of autophagy across subcellular regions of the neuron.

**HIGHLIGHTS and eTOC Blurb:**

- Autophagosome closure, but not maturation, occurs locally at presynaptic sites
- Retrograde transport of autophagosomes requires the motor adaptor UNC-16/JIP3
- The autophagy protease ATG-4.2, but not the related ATG-4.1, is required for autophagosome maturation and degradation
- Defects in retrograde transport and maturation genetically interact and enhance accumulation of autophagosomes in presynaptic regions

## INTRODUCTION

Macroautophagy (also termed autophagy) is a cellular degradation process capable of removing bulk cytoplasm or organelles from cells. Autophagy is well conserved from yeast to mammalian cells, and is best known for its roles in cellular homeostasis (Yin et al., 2016; Zhang and Baehrecke, 2015). Neurons are especially vulnerable to defects in autophagy (Liang and Sigrist, 2017; Son et al., 2012; Vijayan and Verstreken, 2017), as they are post-mitotic cells incapable of diluting defective proteins through repeated cell divisions. Disruption of autophagy is associated with neuronal dysfunction, neurodegeneration and disease (Nah et al., 2015; Nixon, 2013; Wong and Cuervo, 2010).

Neurons are highly polarized cells in which autophagy is spatially compartmentalized. In primary neurons, autophagosomes preferentially form at the distal end of the axon, indicating compartmentalization of autophagosome biogenesis in neurites (Maday et al., 2012). Local formation of autophagosomes is also observed at synapses (Maday and Holzbaur, 2014; Soukup et al., 2016; Stavoe et al., 2016). Synaptic autophagy can be enhanced under conditions of prolonged neuronal activity, suggesting a physiological link between neuronal activity and local autophagosome biogenesis (Shehata et al., 2012; Soukup et al., 2016; Wang et al., 2015). In *Drosophila* neuromuscular junctions, autophagosome biogenesis occurs at presynaptic sites even when motor nerves are severed from the cell bodies, indicating local production of autophagosomes (Soukup et al., 2016). Studies in cultured neurons reveal that local autophagosome biogenesis requires the ordered recruitment of assembly factors to the distal axon (Maday and Holzbaur, 2014; Maday et al., 2012). Similarly, in intact neurons of *C. elegans,* where autophagosomes are observed to form near presynaptic sites, biogenesis depends on the local transport of transmembrane protein ATG-9 to synapses (Stavoe et al., 2016). Together, these studies indicate that autophagy can be locally regulated to result in compartmentalized autophagosome biogenesis in neurons.

Autophagosomes are degraded upon fusion with late endosomes or lysosomes, a process which involves the small GTPase Rab-7 and SNARE machinery (Gutierrez et al., 2004; Hyttinen et al., 2013; Itakura et al., 2012; Jager et al., 2004). In neurons, where lysosomes are preferentially enriched in the cellular soma, autophagosomes in the distal axon are transported towards the soma for degradation (Hollenbeck, 1993; Kaasinen et al., 2008). Retrograde transport of autophagosomes is tightly controlled through both the recruitment and regulation of the dynein motor complex (Ikenaka et al., 2013; Katsumata et al., 2010; Maday et al., 2012). Dynein is recruited to autophagosomes through fusion of autophagosomes with late endosomes (Cheng et al., 2015), and robust retrograde transport is facilitated by the motor scaffolding protein JIP1 (Fu et al., 2014). The importance of these regulated processes in neuronal autophagy is perhaps best exemplified by the consequences of their disruption, including autophagosome accumulation at presynaptic terminals, Alzheimer’s disease-like autophagic stress and axonal pathology (Ikenaka et al., 2013; Lee et al., 2011; Nixon et al., 2005; Takats et al., 2013; Tammineni et al., 2017).

While it is known that autophagy is spatially compartmentalized in neurons, and that regulation of this compartmentalization is important for neuronal physiology, less is known about how the stepwise progression of autophagy maps to the cell biology of the neuron, particularly *in vivo*. Where are the different enzymes of this multi-step process required in the context of the polarized neuron? What mechanisms regulate integration of the distinct steps of autophagy across neuronal regions?

In this study we use forward and reverse genetics, combined with *in vivo* imaging, to systematically dissect the cell biology of autophagy in the neuron. We visualize autophagosomes at single-neuron resolution by examining the localization of the autophagosome-associated protein LGG-1/Atg8/GABARAP (Manil-Segalen et al., 2014; Melendez et al., 2003; Stavoe et al., 2016; Zhang et al., 2015). We find that genes required for autophagosome closure, *atg-2* and *epg-6/WIPI3/4,* are also required for autophagosomes to leave the synapse, but that the autophagosome maturation/acidification gene *epg-5* is not. We identify a new role for the motor adaptor protein, UNC-16/JIP3, in facilitating robust retrograde transport of autophagosomes. To then uncover mechanisms that coordinate transport and clearance of autophagosomes from the neurite, we perform an enhancer screen in *unc-16/jip3* mutant animals and identify a novel dominant enhancer mutation in the poorly understood autophagy gene *atg-4.2*. Mutations in *atg-4.2;unc-16* double mutants display dramatic accumulation of autophagosomes in the neurite, while *atg-4.2* single mutants accumulate autophagosomes in the cell body, phenotypes not observed for the other *atg-4* gene in *C. elegans*, *atg-4.1*. We observe that accumulated autophagosomes in *atg-4.2* mutants fail to mature and degrade, supporting a model in which the cysteine protease *atg-4.2* is specialized to remove LGG-1 from autophagosomes and enable their fusion with lysosomes and degradation. Our studies uncover novel mechanisms that regulate transport and clearance of autophagosomes in neurites and provide a framework to understand how these molecular pathways are coordinated across sites in the neuron *in vivo*.

## RESULTS

### Autophagosomes at *C. elegans* synapses are transported towards and acidified near the cell body

To examine the cell biology of autophagy in neurons of living animals, we imaged LGG-1 in the AIY interneurons of *C. elegans* (schematic, Figure 1A and as previously described in (Stavoe et al., 2016)). LGG-1, a homologue of Atg8 (in yeast) and GABARAP (in mammals), associates with immature and mature autophagosomal structures and is a marker used to track autophagosomes in living cells ((Alberti et al., 2010; Manil-Segalen et al., 2014; Melendez et al., 2003) for validation and controls of the autophagosome marker in AIY interneurons, please see (Stavoe et al., 2016) and Experimental Procedures). Through cell-specific expression in AIY interneurons we visualized autophagosome biogenesis and transport in intact cells and *in vivo* under physiological conditions.

**Figure 1.**
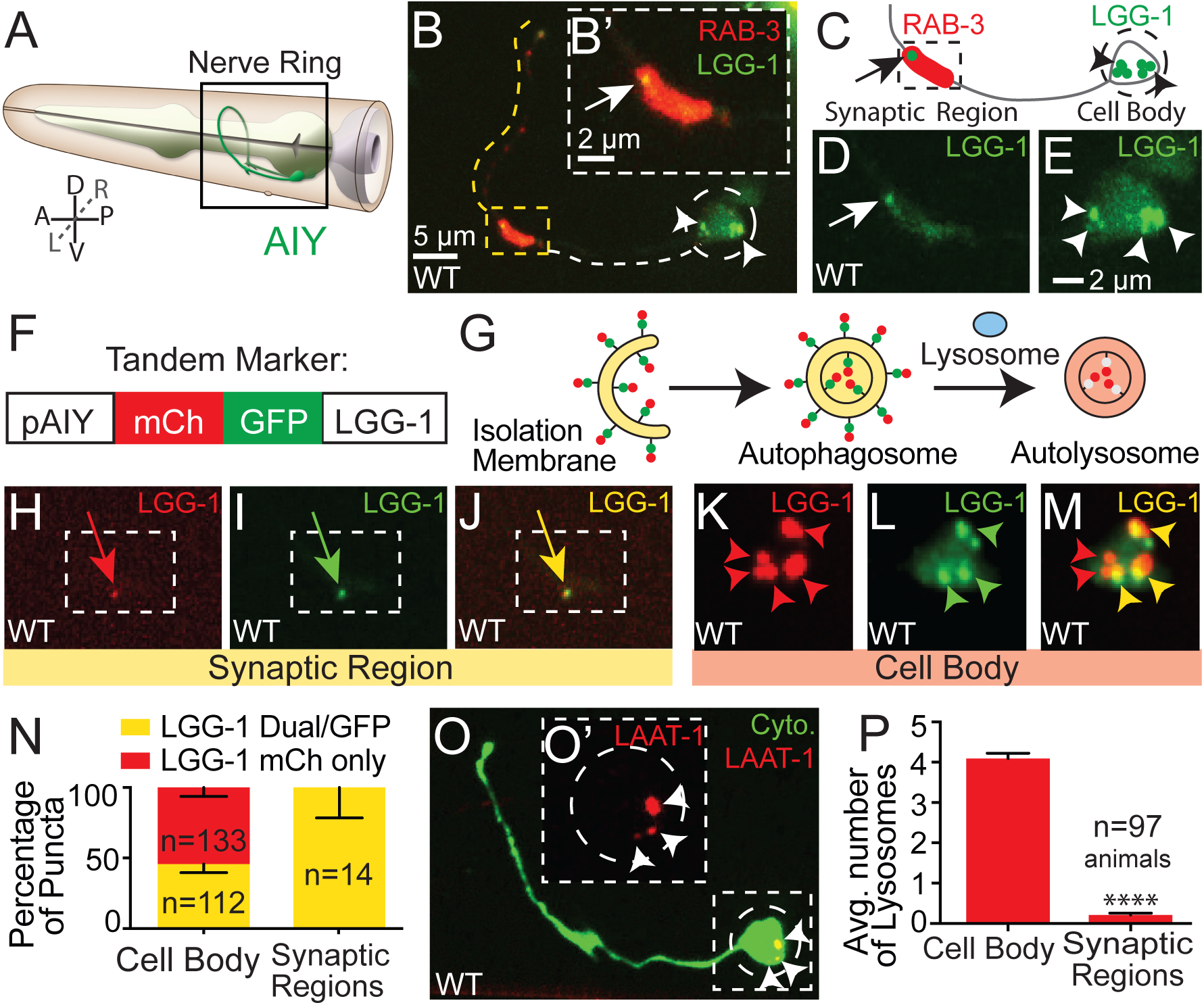
Synaptic autophagosomes are acidified in or near the cell body. (A) Schematic of the AIY interneuron pair (green) located in the nerve ring (boxed) within the head of the worm. Dorsal (D), ventral (V), anterior (A), posterior (P), left (L), right (R). Modified and reprinted with permission from wormatlas.org. (B) Autophagosomes (visualized with GFP::LGG-1) in the synaptic regions (visualized with mCh::RAB-3) of a representative wild type AIY interneuron. The dashed line traces the path of the AIY neurite (see schematic in A). The yellow dotted line highlights the synaptic regions of the neurite, and the white dashed line the asynaptic region. The synapse-rich portion of the synaptic region (boxed) is enlarged in (B’). The cell body is enclosed in a dashed circle. Throughout the figures, images depict maximal projections of a confocal z stacks, arrowheads refer to autophagosomes in the cell body, and arrows point to autophagosomes in the synaptic regions. (C) Schematic of AIY with the cell body and the proximal portion of the synaptic region. (D) As (B’) but GFP::LGG-1 only. Note that we observe autophagosomes in both the cell body and synapse of neurons. LGG-1 puncta numbers are regulated by the autophagy machinery and LGG-1 does not associate with these structures when mutated to prevent lipidation, LGG-1(G116A) (described in Experimental Procedures and (Stavoe et al., 2016)). (E) Enlargement of the cell body in (B) and showing GFP::LGG-1 only. (F and G) Schematic for the tandem LGG-1 marker (mCh::GFP::LGG-1), showing the construct configuration with the AIY promoter (F) and the progression of the tandem LGG-1 marker through the stages of autophagosome biogenesis, fusion with lysosomes and acidification (G and (Chang et al., 2017; Kimura et al., 2007)). (H - M) Autophagosomes in a representative wild type AIY neuron, visualized with the tandem marker (mCh::GFP::LGG-1) in the synaptic region (H-J) and the cell body (K-M). The mCherry channel (H and K), GFP channel (I and L) and merge (J and M) are shown separately. In H-J, the dashed box encloses the synapse-rich portion of the synaptic region, similar to boxed region in B and B’. In J and M, red arrows and arrowheads refer to puncta labeled with only mCh (representing acidified autophagosomes), while yellow arrows and arrowheads refer to structures labeled with both mCh and GFP (representing autophagosomes that have not yet been acidified) (Chang et al., 2017; Kimura et al., 2007). (N) Quantification of puncta labeled with the LGG-1 tandem marker in the cell body and synaptic regions. Error bars show a 95% confidence interval. “n” is the number of puncta assessed counted from 31 neurons. Note that synaptic regions have autophagosomes that are not yet acidified (retain GFP signal), while autophagosomes in the cell body contain both a population of acidified (mCherry only) and immature autophagosomes. (O) The AIY neuron visualized with mCherry (pseudo-colored green) and containing lysosomes, visualized with Lysosome Amino Acid Transporter 1 (LAAT-1)::GFP (Liu et al., 2012) (pseudo-colored red) in the cell body. The dashed circle encloses the cell body, and the dashed box is enlarged in the inset (O’), and shows lysosomes at the cell body. (P) Quantification of the average number of lysosomes in the cell body and synaptic regions. LAAT-1-containing structures are sometimes present in the synaptic regions (28% of examined animals, n=97), but are primarily localized to the cell body in AIY (100% of animals, n=97). Error bars show standard error of the mean (SEM); ****p<0.0001 by Student’s t test. Scale bar in (B) for (B) and (O); in (B’) for (B’), (D), (H)-(J); in (E) for (E), (K)-(M), and (O’).

Autophagosomes in AIY are present in the neurite (Figures 1B-1D) and in the cell body (Figure 1E; (Stavoe et al., 2016)). In the neurite, the majority of autophagosome biogenesis events occur near synaptic regions. We observe that in living animals the number of synaptic autophagosomes in AIY predictably varies depending on the firing state of the neuron, which we can manipulate by altering physiological stimuli that promote AIY responses (Figures S1A-S1C, and S1F), by genetically inhibiting synaptic transmission (Figures S1D, S1E, and S1G) or by chemo-genetically altering the response state of the neuron (Figure S1H). Our findings are consistent with and extend observations that the state of neuronal activity is linked to autophagosome formation at the synapse (Shehata et al., 2012; Soukup et al., 2016; Wang et al., 2015).

Autophagosomes in the distal axon of cultured cells are transported, acidified through fusion with late endosomes or lysosomes and cleared through degradation (Cheng et al., 2015; Maday et al., 2012). Impairment of autophagosome degradation in vertebrate neurons results in accumulation of toxic protein aggregates, axon pathology and neurodegenerative disease (Tammineni et al., 2017). The acidification and degradation of synaptic autophagosomes has not been documented *in vivo*. To better understand the mechanistic regulation of biogenesis, maturation, and degradation of synaptic autophagosomes, we imaged the acidification state of autophagosomes in AIY neurites by employing a tandem label strategy (Chang et al., 2017; Kimura et al., 2007) in which LGG-1 is fused to both mCherry and GFP (Figure 1F). Immature autophagosomes are labeled with both GFP and mCherry, but because GFP is preferentially quenched in an acidic environment, mature structures lose their GFP signal and display solely mCherry signal (schematic, Figure 1G). Using this approach *in vivo* and in single neurons, we observed that 100% (n=14) of synaptic LGG-1 labeled structures retained GFP signal, suggesting that autophagosomes at synapses are not yet mature. In contrast, examination of LGG-1 labeled structures in the asynaptic region (proximal to the cell body) or in the cell body of the neuron demonstrated that over 50% of the structures are positive for mCherry, but not for GFP (57% in asynaptic zone 1 (n=7 puncta); 54% in the cell body (n=245 puncta); from 31 neurons; Figures 1F-1N). Lysosomes promote acidification of autophagosomes, and consistent with synaptic autophagosomes being transported towards the cell body for acidification and degradation, we observe lysosomes preferentially localize to the cell body in AIY (Figures 1O and 1P).

Our findings indicate that while autophagosome biogenesis occurs at the synapse, mature and acidified autolysosomes are preferentially present at or near the cell body. Our *in vivo* observations for synaptic autophagosomes are consistent with studies in cultured vertebrate neurons that demonstrate that fusion of axonal autophagosomes with late endosomes/lysosomes promote retrograde transport, and with studies that demonstrate that autophagosomes are progressively more acidic closer to the cell body (Cheng et al., 2015; Maday et al., 2012). Importantly our *in vivo* system provides a platform to understand the mechanisms that regulate the integration of the distinct steps of autophagy across the neuronal cell.

### Autophagosome closure, but not maturation, is necessary for retrograde transport of synaptic autophagosomes

To map where within the neuron the distinct steps of autophagy are required, we recorded and tracked individual synaptic autophagosomes in wild type and autophagy mutant animals. We found that for wild type animals, autophagosome biogenesis events in the synaptic region average 12 minutes in duration from the first frame of detection until trafficking (with a range of 6 to 23 minutes, n=7). Moreover, all observed LGG-1-containing synaptic structures present at the start of imaging left the synaptic region within 30 minutes (n=12) (Figure 2F). Our findings indicate that following biogenesis at the synapse, autophagosomes are efficiently engaged in retrograde transport towards the cell body.

**Figure 2.**
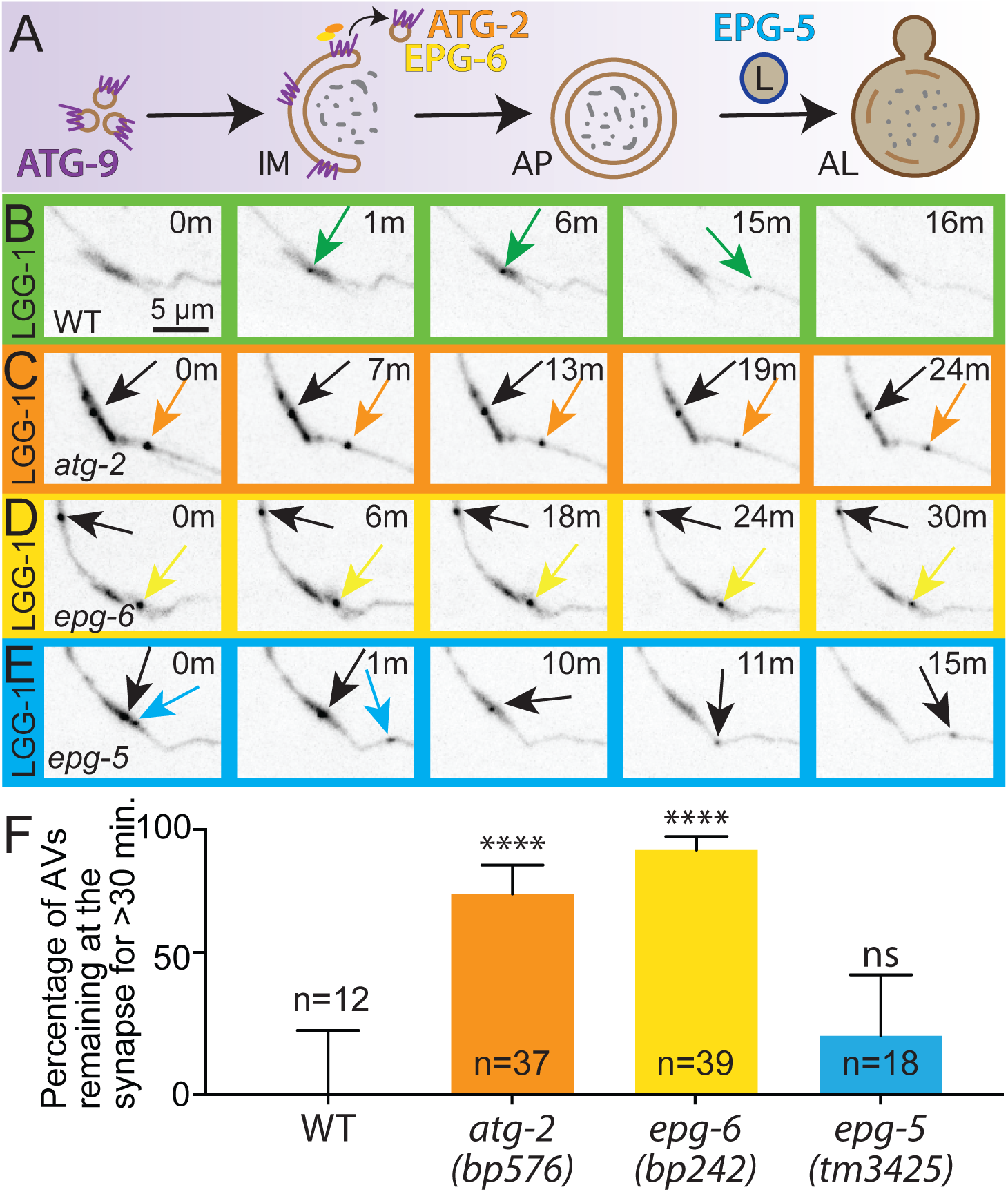
Autophagosome closure, but not acidification, is required to clear autophagosomes from the synaptic region. (A) Schematic of the autophagy pathway highlighting the roles of ATG-2 and EPG-6, required for autophagosomes closure (Lu et al., 2011; Velikkakath et al., 2012), and EPG-5 required for autophagosome maturation (Wang et al., 2016; Zhao et al., 2013). Structure names abbreviated: isolation membrane (IM), autophagosome (AP), lysosome (L) and autolysosome (AL). (B-E) Time series showing dynamics of autophagosomes (visualized with GFP::LGG-1) in the synaptic-rich zone 2 region of representative wild type (B) and autophagy mutant animals *atg-2(bp576)* (C), *epg-6(bp242)* (D), and *epg-5(tm3425)* (E). Within each time series, uniquely colored arrows follow individual autophagosomes over time. Time point unit “m” is minutes. While images bleach with prolonged imaging, note how autophagosomes in *atg-2* and *epg-6* mutants remain stable at the synapse during the time course, while in wild type and *epg-5* mutant animals autophagosomes traffic back towards the cell body. See also Supplemental Movies S1-S4. (F) Quantification of the percentage of autophagosomes that are present at the start of imaging and remain at the synapse after 30 minutes. ****p<0.0001 by Fisher’s exact test between wild type and mutant animals. Error bars show a 95% confidence interval. Abbreviations: “ns” is not significant as compared to wild type; “n” is number of puncta assessed; “AV” is autophagic vacuole. Quantifications were performed on maximal projections of confocal z-stacks acquired using identical imaging conditions across genotypes. Scale bar in (B) for (B)-(E).

We then examined if later steps of autophagy, such as autophagosome maturation, were necessary for retrograde transport. ATG-2 and EPG-6/WIP3/4 are necessary for autophagosome closure (Lu et al., 2011; Velikkakath et al., 2012), while EPG-5 is required for autophagosome maturation (Wang et al., 2016; Zhao et al., 2013) (schematic in Figure 2A). We observe that, unlike wild type animals, 76% of LGG-1-containing structures in *atg-2* mutants (n=37) and 92% in *epg-6* mutants (n=39) remained at the synapse after 30 minutes (Figures 2C, 2D, 2F and Supplemental Movies S1-S3), suggesting that completion of autophagosome closure is necessary for initiation of retrograde transport. However, *epg-5* mutant animals, in which autophagosomes are able to close but not be acidified (Zhao et al., 2013), phenocopied wild type animals (n=18) (Figures 2E, 2F and Supplemental Movie S4). Our findings indicate that while autophagosome closure is necessary for efficient retrograde transport of synaptic autophagosomes, autophagosome maturation is not. Our genetic findings are consistent with our cell biological observations that maturation of autophagosomes occurs near the neuronal cell body, and support the model that different steps of autophagy occur at distinct subcellular compartments in neurons. Our findings also extend our understanding of the differential requirement of these late steps of autophagy in the context of the cell biology of the neuron, and link regulated closure of autophagosomes with retrograde transport, maturation and removal.

### Dynactin and JIP3/UNC-16 promote autophagosome retrograde transport and retention in the cell body

To investigate the removal of synaptic autophagosomes, we first examined the molecular mechanisms underlying the trafficking of autophagosomes from the synapse *in vivo*. In *C. elegans*, as in vertebrate neurons, retrograde transport of autophagosomes depends on the dynein complex (Cheng et al., 2015; Ikenaka et al., 2013; Katsumata et al., 2010; Maday et al., 2012) (Figures 3A-3C). We therefore examined known regulators of transport to investigate their specific requirement in autophagosome trafficking.

**Figure 3.**
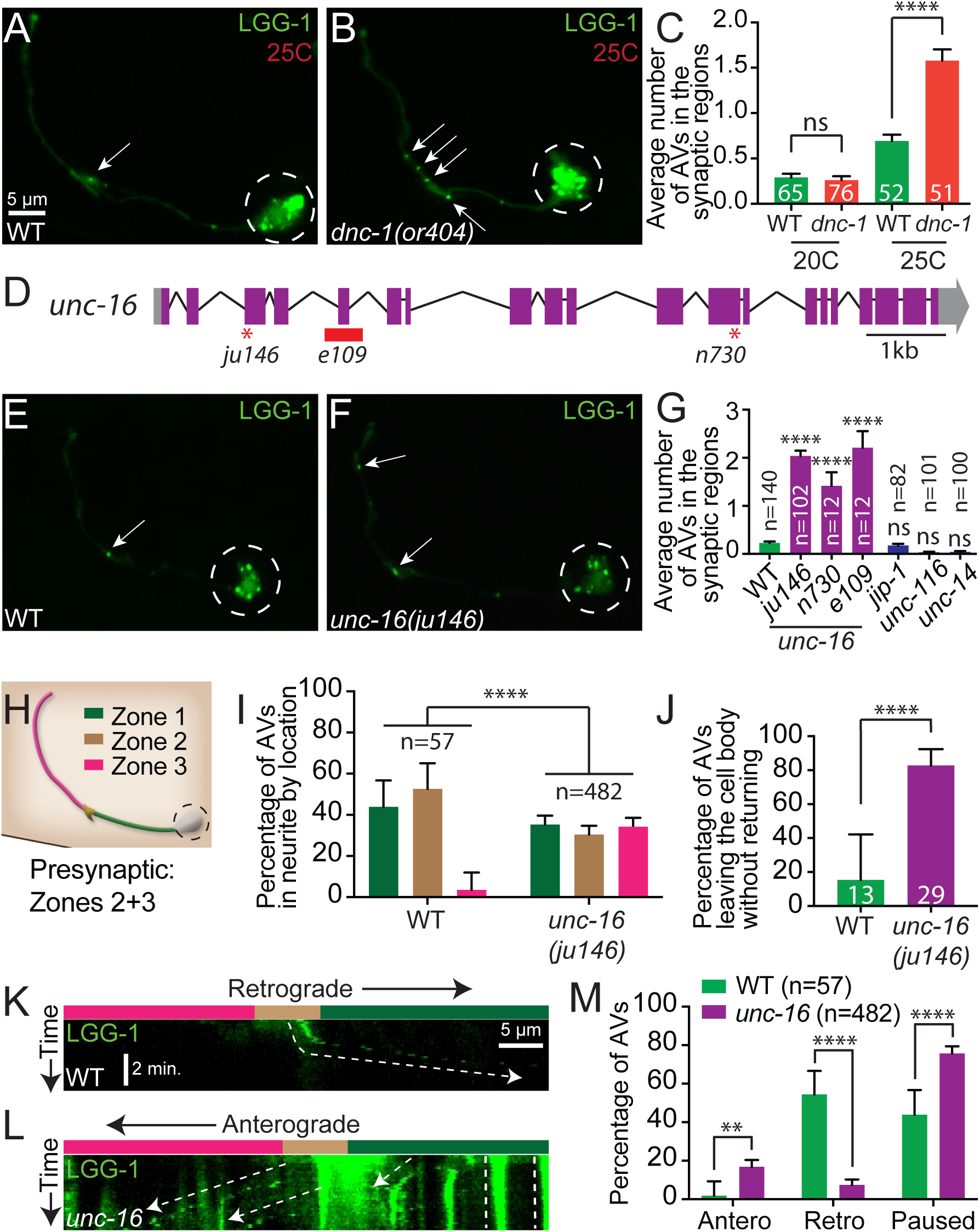
Dynactin and UNC-16/JIP3 mediate retrograde transport of autophagosomes from the synaptic regions. (A and B) Autophagosomes (visualized with GFP::LGG-1) in a wild type AIY neuron (A) and in a temperature-sensitive dynactin, *dnc-1(or404)* mutant animal (B) grown for 48 hours at the restrictive temperature of 25C. Arrows denote autophagosomes in the synaptic region. Dashed circles enclose the cell bodies. (C) Quantification of the average number of autophagosomes in the synaptic regions in wild type and *dnc-1(or404)* mutant animals kept for 48 hours at the permissive (20C) or restrictive (25C) temperature. Error bars show standard error of the mean (SEM). ****p<0.0001 by one-way ANOVA with Tukey’s post hoc analysis between indicated groups. Abbreviations: “ns” is not significant; “AV” is autophagic vacuole. Numbers on bars represent number of animals examined. (D) Schematic of the *unc-16* genomic region with exons (purple boxes), untranslated regions (grey boxes), introns (black lines) and allelic lesions (early stop codons as red asterisks in *ju146* and *n730*; and deletion as red box in *e109*). Scale bar shows a 1 kilobase (kb) region. (E and F) Autophagosomes (visualized with GFP::LGG-1) in a representative wild type AIY neuron (E) and in an *unc-16(ju146)* mutant (F) grown at the standard 20C. Arrows denote autophagosomes in the synaptic region. Dashed circles enclose the cell bodies. (G) Quantification of the average number of autophagosomes in the synaptic regions in wild type, three alleles of unc-16 (*ju146*, *n730*, and *e109*), *jip-1(gk466982)*, *unc-116(e2310)*, and *unc-14(e57)* mutant animals. Error bars show standard error of the mean (SEM). ****p<0.0001 by one-way ANOVA with Tukey’s post hoc analysis. Abbreviations: “n” is the number of animals; “ns” is not significant as compared to wild type; “AV” is autophagic vacuole. (H) Schematic of the AIY interneuron colored by region and similar to the schematic in Figure 1A: Zone 1 (asynaptic, green), Zone 2 (enriched synaptic region, brown), and Zone 3 (punctate synaptic region, pink, see (Colon-Ramos et al., 2007)). A dashed circle encloses the cell body. Modified and reprinted with permission from wormatlas.org. (I) Quantification of the percentage of autophagosomes in the AIY neurite by location. Bars are colored as zones in (H). Zone-1 localized autophagosomes observed in wild type animals many times represent synaptic autophagosomes in transit towards the cell soma. Error bars show a 95% confidence interval. ****p<0.0001 by Chi-square test between wild type (n=57 autophagosomes from 36 animals) and *unc-16* mutants (n=482 autophagosomes from 37 animals). Quantifications were performed on maximal projections of confocal z-stacks acquired using identical imaging conditions across genotypes. (J) Quantification of the percentage of autophagosomes that traffic out of the cell body and do not return within the 5-minute imaging window in wild type (n=13 autophagosomes) and *unc-16* mutants (n=29 autophagosomes). Error bars show a 95% confidence interval. ****p<0.0001 by Fisher’s exact test between wild type and mutants. (K and L) Kymographs of autophagosome trafficking in AIY neurites in representative wild type (K) and *unc-16* mutant (L) animals showing retrograde (right down diagonals), anterograde (left down diagonals), and paused events (vertical lines). Neurite zones are indicated by color above kymographs and as in (H). (M) Quantification of autophagosome trafficking events in the AIY neurite, showing percentage of autophagosomes that undergo anterograde, retrograde or paused events within the 5-minute imaging window for wild type (green; n=57 autophagosomes in 71 neurons) and *unc-16* mutant animals (purple; n=482 autophagosomes in 73 neurons). ****p<0.0001, **p<0.01 by Fisher’s exact test between wild type and mutant animals. Error bars show a 95% confidence interval. Scale bar(s) in (A) for (A), (B), (E) and (F); in (K) for (K) and (L).

We observed that while *jip-1/Mapk8Ip1*, *unc-116/kinesin-1* and *unc-14/RUN* mutants did not display defects in autophagosome accumulation at synapses (Figure 3G), loss of JIP3/UNC-16/Sunday Driver resulted in a significant accumulation of autophagosomes in the neurite, but not in the cell body (Figures 3E-3G, and 6G). The *unc-16/jip3* mutant phenotype was similar to the phenotype seen for *dnc-1/p150* dynactin complex subunit mutants (which affect the dynein complex) ((Ikenaka et al., 2013) and Figures 3B-3C). JIP3/UNC-16/Sunday Driver is a conserved adaptor protein that binds to kinesin and dynein (Arimoto et al., 2011; Cavalli et al., 2005) to regulate early endosome and lysosome transport (Brown et al., 2009; Byrd et al., 2001; Drerup and Nechiporuk, 2013; Edwards et al., 2015; Edwards et al., 2013; Gowrishankar et al., 2017). Three independent alleles of *unc-16* (*ju146, e109,* and *n730*) (Figure 3D) result in an increase from 34% (in wild type) to 100% (in *unc-16* alleles) of animals with autophagosomes in the neurites. Moreover, while wild type animals average less than one autophagosome per neurite, *unc-16* mutants average ∼2 autophagosomes in the presynaptic regions (Figure 3G), indicating abnormal accumulation of autophagosomes. Analyses of the subcellular localization of LGG-1-containing structures in *unc-16/jip3* mutants also revealed a higher probability for autophagosomes to be present in the distal synaptic Zone 3 region (Figures 3H and 3I). The distal accumulation of autophagosomes observed for *unc-16* mutants was not due to a rearrangement of microtubule polarity, as we observed that the microtubule plus-end-binding protein EBP-2/EB1 (Baas and Lin, 2011; Maniar et al., 2011) is similarly oriented (plus-end-out) in wild type and *unc-16/jip3* mutant animals (Figure S2). Our findings indicate that *unc-16/jip3* is necessary to prevent the abnormal accumulation of autophagosomes in the neurite, and are consistent with a requirement for *unc-16/jip3* in autophagosome retrograde transport.

Previous reports have implicated UNC-16/JIP3/Sunday Driver in the regulation of retrograde transport and retention of lysosomes, early endosomes and Golgi in the cell body ((Brown et al., 2009; Byrd et al., 2001; Edwards et al., 2013) and Figures S3). The role of UNC-16/JIP3 in transport of autophagosomes has not been examined. To directly investigate if *unc-16/jip3* is required for retrograde transport of autophagosomes, we performed time-lapse imaging of individual autophagosomes in wild type neurons (n=71) and *unc-16* mutant neurons (n=73). We observed that in wild type animals 54% of autophagosomes in neurites (n=57 autophagosomes) traffic in the retrograde direction towards the cell body, while in *unc-16/jip3* mutants only 7% (n=482 autophagosomes) traffic in the retrograde direction (Figures 3K-3M). Instead *unc-16/jip3* mutant animals display increased anterograde trafficking of autophagosomes (from 2% in wild type to 17% in *unc-16* mutants) and increased number of paused autophagosomes (defined as autophagosomes that do not traffic within a 5 minute observation window; from 44% in wild type to 76% in *unc-16* mutants) (Figures 3K-3M). In wild type animals, autophagosomes in the cell body are sometimes transported in an anterograde fashion into the neurite, but 85% of those autophagosomes (n=13 autophagosomes) return to the cell body via retrograde transport within the 5-minute examination window. In contrast, in *unc-16* mutant neurites only 17% of autophagosomes (n=29 autophagosomes) that leave the cell body return within 5 minutes (Figure 3J). Our findings are consistent with previous reports that demonstrate that another JIP family member (JIP1) is important for regulating autophagosome retrograde transport in cultured vertebrate neurons (Fu et al., 2014). We now extend those findings and demonstrate an important role for *unc-16/jip3* in regulating the retrograde trafficking of autophagosomes *in vivo*.

To examine how UNC-16-dependent retrograde transport contributes to the acidification and degradation of autophagosomes, we then tested if defects in *unc-16* mutants altered the total number of autophagosomes in neurons. We observed that the total number of autophagosomes in neurons of *unc-16/jip3* mutants is higher than seen in wild type animals (average from 4.0 in wild type neurons (n=65) to 6.4 in *unc-16* mutant neurons (n=64)). Our findings indicate that the increased number of autophagosomes observed in the neurites of *unc-16/jip3* mutants does not result from a simple redistribution of a fixed number of autophagosomes. Instead, retrograde transport, promoted by *unc-16/jip3*, appears to be necessary for the effective degradation of autophagosomes, and defects in transport in *unc-16/jip3* mutants therefore result in an abnormal accumulation of autophagosomes. Our findings underscore the importance of regulated trafficking in the degradation of synaptic autophagosomes.

### Mutant allele *ola316* enhances the number of autophagosomes in the neurites of *unc-16/jip3* mutants

Degradation of autophagosomes occurs upon fusion with lysosomes, an important and final step that is tightly regulated but poorly understood (Itakura et al., 2012; Nakamura and Yoshimori, 2017). To better understand the molecular machinery underlying coordinated trafficking and degradation of synaptic autophagosomes, we performed a visual forward genetic screen in the *unc-16/jip3* mutant background. We reasoned that molecules important for regulated lysosomal fusion will result in an enhanced accumulation of autophagosomes, and screened for mutants with increased numbers of autophagosomes in the AIY synaptic regions. We uncovered a mutant allele, *ola316*, which displays a dramatic accumulation of autophagosomes in the neurites. While *unc-16/jip3* mutants display an average of 2.6 autophagosomes abnormally accumulated in the synaptic regions (n=64 neurites), *ola316;unc-16* double mutants enhance this phenotype and display an average of 9.4 autophagosomes in the synaptic regions (n=67 neurites) (Figures 4A-4D).

**Figure 4.**
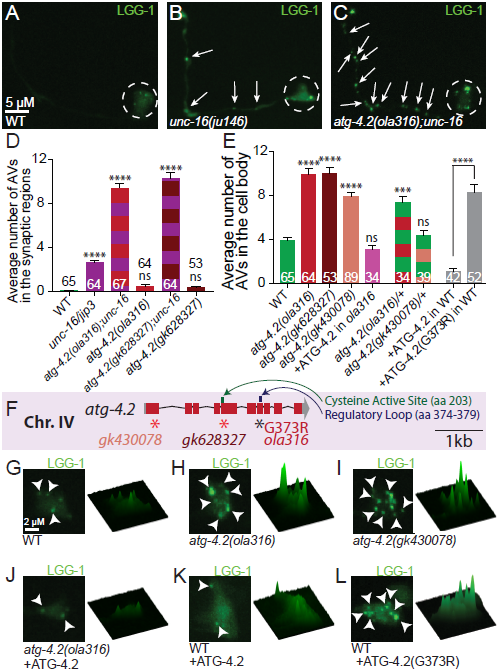
ATG-4.2 enhances UNC-16/JIP3-mediated accumulation of autophagosomes. (A-C) Autophagosomes (visualized with GFP::LGG-1) in representative wild type (A), *unc-16(ju146)* mutant (B), and *atg-4.2(ola316);unc-16(ju146)* double mutant (C) AIY neurons. Arrows denote autophagosomes in the neurite. Dashed circles enclose the cell bodies. (D) Quantification of the average number of autophagosomes in the synaptic regions (Zones 2+3) of the AIY neurite. Synaptic quantifications for wild type (green bar), *unc-16/jip3* mutant (purple bar), *atg-4.2(ola316);unc-16* double mutant (purple and red striped bar), *atg-4.2(ola316)* mutant (red bar), *atg-4.2(gk628327);unc-16* mutant (purple and maroon striped bar), and *atg-4.2(gk628327)* mutant (maroon bar) animals are shown. Error bars show standard error of the mean (SEM). ****p<0.0001 by one-way ANOVA with Tukey’s post hoc analysis between wild type and mutants. The numbers on the bars represent number of neurons examined. Abbreviations: “ns” is not significant as compared to wild type; “AV” is autophagic vacuole. Scoring was done by blindly quantifying maximal projections of confocal z-stacks. (E) Quantification of the average number of autophagosomes in the cell body of AIY in wild type and mutant animals. Cell body quantifications for wild type (green bar), *atg-4.2(ola316)* mutant (red bar), *atg-4.2(gk628327)* mutant (maroon bar), *atg-4.2(gk430078)* mutant (peach bar), cell-specific expression of an ATG-4.2 rescuing array in *atg-4.2(ola316)* mutants (pink bar), heterozygotes for *atg-4.2(ola316)* (red and green striped bar), heterozygotes for *atg-4.2(gk430078)* (peach and green striped bar), cell-specific expression of wild type ATG-4.2 or ATG-4.2 cDNA containing the lesion found in allele *atg-4.2(ola316)* (corresponding to (G373R)) and expressed in wild type animals (grey bars). Note that *atg-4.2(ola316)* behaves as a dominant negative lesion in that animals heterozygous for the *atg-4.2(ola316)* allele but not for a putative null allele, *atg-4.2(gk430078)*, displayed the phenotype, and in that overexpression of mutant ATG-4.2(G373R), but not wild type ATG-4.2, induced the phenotype (grey bars). Error bars show standard error of the mean (SEM). ***p<0.001; ****p<0.0001 by one-way ANOVA with Tukey’s post hoc analysis between wild type and mutants or as indicated. The numbers on the bars represent number of neurons examined. Abbreviations: “ns” is not significant as compared to wild type; “AV” is autophagic vacuole. Scoring was done by blindly quantifying maximal projections of confocal z-stacks. (F) Schematic of the *atg-4.2* genomic region on chromosome IV with exons (red boxes), untranslated regions (grey boxes), and introns (black lines). Alleles affect indicated regions with early stop codons (red asterisks; *gk430078* and *gk628327*) or amino acid change (G373R for *ola316*). The crystal structure of the human Atg4B-LC3 complex revealed a regulatory loop important for LC3 recognition by Atg4B (Kumanomidou et al., 2006). Alignment with *C. elegans atg-4* homologs showed that this regulatory loop (dark blue, amino acids 374-379) is conserved (Wu et al., 2012). The G373R lesion in atg-4.2(*ola316)* neighbors the regulatory loop, and is also sterically close to the cysteine active site (dark green, amino acid 203) in the three dimensional predicated structure of the protein (Wu et al., 2012), which might explain why it behaves as a dominant negative lesion. Scale bar shows a 1 kilobase region. (G-L) Autophagosomes in the AIY cell body imaged as maximal projection of confocal z stacks (left) and corresponding surface plots of the LGG-1 signal intensity (right) in wild type (G); *atg-4.2(ola316)* mutants (H); *atg-4.2(gk430078)* mutants (I); *atg-4.2(ola316)* mutants containing an ATG-4.2 rescue array (J); wild type cell-specifically overexpressing ATG-4.2 (K); or ATG-4.2(G373R) (L). Arrowheads denote autophagosomes in the cell body. Scale bar in (A) for (A)-(C); in (G) for (G)-(L).

We then examined if the *ola316* allele displayed a phenotype independent of the *unc-16/jip3* mutant lesion. To achieve, this we outcrossed the *unc-16/jip3* lesion and examined LGG-1 containing structures in the *ola316* single mutants. We observed that, unlike the *ola316;unc-16* double mutants, the *ola316* single mutants did not display a significant accumulation of autophagosomes in the synaptic regions when compared with *unc-16* or *unc-16;ola316* double mutants (n= 64 neurites) (Figure 4D). However, we did observe an increased number of autophagosomes in the cell body of the *ola316* single mutants, from an average of 3.9 autophagosomes in wild type (n=65) to 9.9 autophagosomes in ola316 mutants (n=64) (Figures 4E, 4G and 4H). We also observed an increase in the percentage of neurites with one or more autophagosomes in the synaptic regions, from 7% in wild type (n=68) to 58% in *ola316* mutant neurites (n=78) (Figure S4A). The gene affected in the *ola316* allele is important for most neurons in the nematode’s nervous system, as abnormal panneuronal accumulation of autophagosomes is observed for both the single mutants and the *ola316;unc-16* double mutants (Figure S5). Together our data indicate that the *ola316* allele affects a gene that is important for preventing the abnormal accumulation of autophagosomes both in the neuronal soma and, through cooperation with UNC-16/JIP3, in the neurites.

### *ola316* is a dominant allele of the autophagy gene *atg-4.2*

Genetic characterization of the *ola316* allele revealed that the phenotype results from a single genetic dominant lesion (Figure 4E, green and red striped bar). To identify the causative lesion in *ola316*, we performed single-nucleotide polymorphism (SNP) mapping, whole genome sequencing, independent allele analyses and rescue experiments. We SNP-mapped *ola316* to a 2.2 Megabase region between 10.1 and 12.3 Mb on chromosome IV. We then performed whole genome sequencing (Minevich et al., 2012; Sarin et al., 2008) and uncovered a missense lesion in exon 7 in the autophagy gene *atg-4.2*. Two independent alleles of *atg-4.2 (gk430078* and *gk628327)*, which result in early stop codons (Figure 4F), phenocopy the *ola316* single mutant phenotype (Figure 4E (maroon and peach bars) and 4I). Moreover *unc-16;atg-4.2(gk628327)* double mutants also phenocopy the *unc-16;ola316* double mutants isolated from the genetic screen (Figure 4D). Consistent with *ola316* being an allele of *atg-4.2*, AIY-specific overexpression of the wildtype *atg-4.2* cDNA in *ola316* mutant animals rescues the *ola316* mutant phenotype (Figures 4E (pink bar), 4H and 4J), while overexpression of mutant *atg-4.2(ola316)* cDNA in wild type animals is sufficient to induce the phenotype (Figures 4E (grey bars), 4K and 4L). Together our data indicate that *ola316* is a dominant allele of the autophagy gene *atg-4.2*, and that ATG-4.2 acts cell autonomously in neurons to regulate the number of LGG-1 structures that otherwise accumulate in the cell body.

Atg4s are a conserved family of cysteine proteases that bind to and cleave Atg8/LC3/LGG-1 proteins to achieve two functions: 1) prime LGG-1 for conjugation onto the autophagosomal membrane; and 2) remove LGG-1 from the autophagosomal membrane (Kirisako et al., 2000; Maruyama and Noda, 2017). Conjugation of Atg8/LC3/LGG-1 to the membrane is necessary for autophagosome elongation and closure (Fujita et al., 2008; Kirisako et al., 1999; Manil-Segalen et al., 2014; Nakatogawa et al., 2007; Weidberg et al., 2010), and removal of Atg8/LC3/LGG-1 is thought to occur prior to degradation (Betin et al., 2013; Kimura et al., 2007; Yu et al., 2012). Therefore, the seemingly opposite roles for Atg4 protease activity on LGG-1 need to be coordinated at different stages of the autophagosome biogenesis and degradation process. How this duality of roles is achieved is not well understood.

In yeast one Atg4 gene performs both the priming and deconjugation events on Atg8/LGG-1 (Kirisako et al., 2000; Maruyama and Noda, 2017), while in higher metazoans multiple genes encode distinct Atg4 cysteine proteases, some with unknown function, specificity, or redundancy (Kauffman et al., 2018; Li et al., 2011; Marino et al., 2003; Wu et al., 2012; Zhang et al., 2016). Consistent with this, the two *C. elegans atg-4* genes (*atg-4.1* and *atg-4.2*) (Figure 5A) were shown to be largely redundant (Wu et al., 2012), with ATG-4.1 displaying enhanced proteolytic activity of a soluble pro-form of LGG-1/Atg8 as compared to ATG-4.2. The specific roles of ATG-4.2 in autophagy, if any, remain unknown.

**Figure 5.**
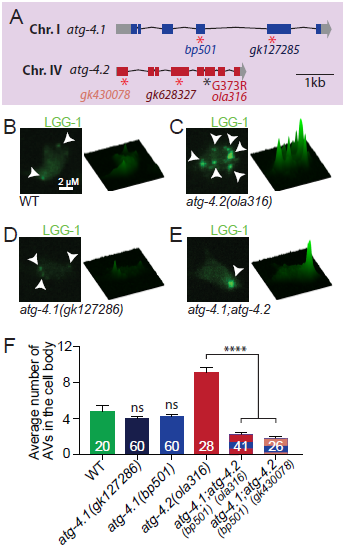
ATG-4.1 and ATG-4.2 have distinct phenotypes in autophagy. (A) Schematic of the *atg-4.1* and *atg-4.2* genomic regions on chromosomes I and IV respectively, with exons (blue or red boxes), untranslated regions (grey boxes), and introns (black lines). Alleles affect indicated regions with early stop codons (red asterisks; *bp501*, *gk127285* in *atg-4.1* and *gk430078* and *gk628327* in *atg-4.2*) or amino acid changes (G373R for *ola316* in *atg-4.2*). Scale bar shows a 1 kilobase region. (B-E) Autophagosomes in the AIY cell body imaged as maximal projection of confocal z stacks (left) and corresponding surface plots of the LGG-1 signal intensity (right) in wild type (B), *atg-4.2(ola316)* mutants (C), *atg-4.1(gk127286)* mutants (D), and *atg-4.1;atg-4.2* double mutants (E). Arrowheads denote autophagosomes in the cell body. Scale bar in (B) for (B)-(E). (F) Quantification of the average number of autophagosomes in the cell body of AIY in wild type and mutant animals. Cell body quantifications for wild type (green bar), *atg-4.1* mutant alleles (navy and blue bars), *atg-4.2* mutants (red bar), and indicated *atg-4.1;atg-4.2* double mutants (red and blue or peach and blue striped bars) are shown. Error bars show standard error of the mean (SEM). ****p<0.0001 by one-way ANOVA with Tukey’s post hoc analysis between wild type and mutants or as indicated. The numbers on the bars represent number of neurons examined; corresponding to ≥10 animals for each genotype. Abbreviations: “ns” is not significant as compared to wild type; “AV” is autophagic vacuole. Scoring was done by blindly quantifying maximal projections of confocal z-stacks.

### ATG-4.2, but not ATG-4.1, is required to prevent autophagosome accumulation

The two *C. elegans* Atg4 homologs display 44% amino acid similarity, including conserved catalytic and regulatory sites (Wu et al., 2012). A previous study investigating the degradation of protein aggregates in embryos found that *atg-4.1* is more efficient at the first LGG-1 cleavage event (priming), and more effective in removal of protein aggregates, than *atg-4.2* (Wu et al., 2012). To examine if *atg-4.1* also acts as *atg-4.2* to prevent autophagosomal accumulation we examined two independent alleles of *atg-4.1*, including a confirmed null allele *(bp501* (null) (Wu et al., 2012) and *gk127286* (early stop codon)). Surprisingly we did not detect LGG-1 puncta accumulation in the examined *atg-4.1* mutant animals (Figures 5D and 5F). Consistent with *atg-4.1* having a distinct phenotype compared to *atg-4.*2, we observed that *atg-4.1* did not enhance the *unc-16* mutant phenotype (Figure 6H). Our findings indicate that *atg-4.1* and *atg-4.*2 have distinct functions *in vivo* that result in different phenotypes regarding the accumulation of autophagosomes.

**Figure 6.**
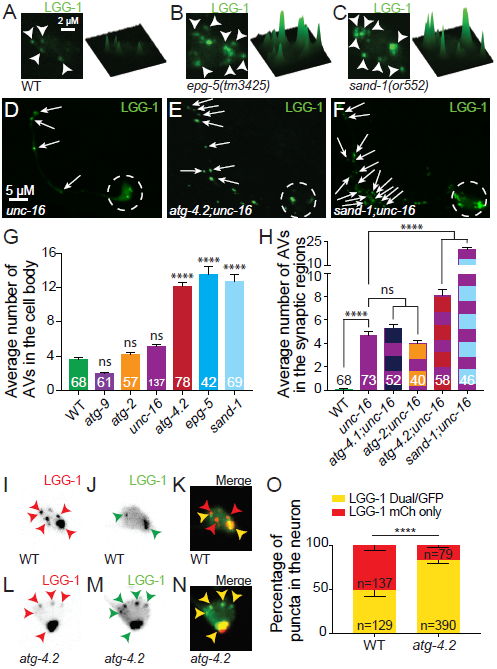
ATG-4.2 promotes autophagosome maturation. (A-C) Autophagosomes in the AIY cell body imaged as maximal projections of confocal z stacks (left) and corresponding surface plots of the LGG-1 signal intensity (right) in representative wild type (A), *epg-5(tm3425)* mutant (B) and *sand-1(or552)* mutant (C) animals. Arrowheads denote autophagosomes in the cell body. (D-F) Autophagosomes (visualized with GFP::LGG-1) in a representative *unc-16(ju146)* mutant (D), *atg-4.2(ola316);unc-16(ju146)* double mutant (E) and *sand-1(or552);unc-16(ju146)* double (F) mutants neurons. Arrows denote autophagosomes in the synaptic regions. The white dashed circles enclose the cell bodies. (G) Quantification of the average number of autophagosomes in the cell body of AIY in wild type and mutant animals. Cell body quantifications for wild type (green bar), early autophagy mutants *atg-9(wy56)* (purple bar), and *atg-2(bp576)* (orange bar), *unc-16(ju146)* mutants (purple bar), *atg-4.2(ola316)* mutants (red bar), and in mutants involved in the later steps of autophagy, *epg-5(tm3425)* (blue bar) and *sand-1(or552)* (light blue bar). Error bars show standard error of the mean (SEM). ****p<0.0001 by one-way ANOVA with Tukey’s post hoc analysis between wild type and mutants. The numbers on the bars represent number of neurons examined. Abbreviations: “ns” is not significant as compared to wild type; “AV” is autophagic vacuole. Scoring was done by blindly quantifying maximal projections of confocal z-stacks. (H) Quantification of the average number of autophagosomes in the synaptic region of AIY in *unc-16* mutant and double mutant animals. Synaptic region quantifications for wild type (green bar), *unc-16(ju146)* mutants (purple bar), *atg-4.1(gk127286)*;*unc-16(ju146)* double mutants (purple and navy striped bar), *atg-2(bp576)*;*unc-16(ju146)* double mutants (purple and orange striped bar), *atg-4.2(ola316)*;*unc-16(ju146)* double mutants (purple and red striped bar), and *sand-1(or552)*;*unc-16(ju146)* double mutants (purple and light blue striped bar) are shown. Note that the increased severity of the *sand-1* enhancement might suggest a more complete block in autophagosome degradation than that of *atg-4.2*, which is partially redundant with *atg-4.1.* The lack of enhancement in the *atg-2;unc-16* and the *atg-4.1;unc-16* double mutants suggests that enhancement is specific to late mutants that block autophagic maturation, and not a general result of autophagy mutants. Error bars show standard error of the mean (SEM). ***p<0.001; ****p<0.0001 by one-way ANOVA with Tukey’s post hoc analysis between wild type and mutants. The numbers on the bars represent number of neurons examined. Abbreviations: “ns” is not significant as compared to *unc-16(ju146)* single mutants; “AV” is autophagic vacuole. Quantifications were blindly scored from maximal projections of confocal z-stacks. (I-N) Autophagosomes in representative wild type (I-K) and *atg-4.2* mutant (L-N) cell bodies, visualized with the tandem marker (mCh::GFP::LGG-1). The mCherry channel (I and L), GFP channel (J and M) and merge (K and N) are shown separately. In the merge images K and N, red arrowheads refer to puncta labeled with only mCh, while yellow arrowheads refer to structures labeled with both mCh and GFP. (O) Quantification of puncta labeled with the LGG-1 tandem marker in wild type and *atg-4.2* mutant animals. Error bars show a 95% confidence interval. ****p<0.0001 by Chi-square test between wild type and mutants. “n” is the number of puncta assessed. For wild type, a different analysis of the same dataset is shown in Figure 1N. Scale bar in (A) for (A)-(C), (I)-(N); in (D) for (D)-(F).

While we observe distinct phenotypes for *atg-4.1* and *atg-4.2* in the accumulation of autophagic structures, we also note that both *atg-4.1* and *atg-4.2* single mutant animals contain LGG-1 puncta in the neuron (Figures 5B-5D and 5F). This observation suggests that either protease is able to partially compensate for the other and perform the first priming cleavage necessary to conjugate LGG-1 onto the nascent autophagosome. Consistent with these two proteases performing partially redundant roles, we have previously shown that in AIY, where autophagy is also required for synapse development, the synaptic assembly phenotype is not seen for *atg-4.1* or *atg-4.2* single mutants, but is evident in the *atg-4.1*;*atg-4.2* double mutants (Figure S6 and (Stavoe et al., 2016)). To further explore the partial redundancy of *atg-4.1* and *atg-4.2* in the accumulation of LGG-1 containing structures, we examined GFP::LGG-1 in the *atg-4.1;atg-4.2* double mutants. We observed a reduction in the number of accumulated LGG-1 puncta for the *atg-4.1;atg-4.2* double mutants as compared to either single mutant (Figures 5E and 5F). Our result indicates that *atg-4.1* and *atg-4.2* act in a partially redundant fashion, that *atg-4.1* activity is necessary to achieve the autophagosome accumulation seen in *atg-4.2* mutants, and that *atg-4.2* activity is necessary in the *atg-4.1* mutants to see normal levels of autophagosomes. Importantly, our findings indicate that *atg-4.1* and *atg-4.2* display distinct phenotypes in LGG-1 puncta accumulation, and that *atg-4.2* is specifically necessary to prevent abnormal accumulation of autophagosomes in the neuronal soma.

### ATG-4.2 promotes autophagosome maturation and removal from the cell body

Our observations that lesions in *atg-4.2* result in an abnormal accumulation of autophagosomes in the neuronal cell body are consistent with a model where *atg-4.2* acts late in the autophagy pathway to promote autophagosome maturation and degradation. Autophagosome maturation into an acidic autolysosome is mediated by fusion between autophagosomes and late endosomes or lysosomes in a process that requires the small GTPase Rab7 (Gutierrez et al., 2004; Hyttinen et al., 2013; Jager et al., 2004). We reasoned that if *atg-4.2* mutants have a defect in lysosomal fusion, then loss of *rab-7* function should phenocopy the *atg-4.2* mutants. Since null lesions in *rab-7* are lethal, we examined mutants for the *rab-7* activator *sand-1/Mon1* (Hegedus et al., 2016; Poteryaev et al., 2007), and the *rab-7* effector, *epg-5* (Wang et al., 2016). We observe that indeed *epg-5* and *sand-1* mutants display increased autophagosome accumulation in the cell body (averaging 13.6 autophagosomes in *epg-5* mutants (n=42 neurons) and 12.7 autophagosomes in *sand-1* mutants (n=69 neurons), compared to 12.1 autophagosomes in *atg-4.2(ola316)* mutants (n=78 neurons) and 3.6 autophagosomes in wild type animals (n=68 neurons); Figures 6A-6C, 6G). The observed phenotypes are specific to late blocks in autophagy, as mutants for genes involved in early events prior to autophagosome closure or trafficking, such as *atg-9*, *atg-2,* or *unc-16* mutants, did not result in autophagosome accumulation in the cell body (Figure 6G). Moreover, *sand-1;unc-16*, like *atg-4.2;unc-16*, also dramatically enhanced the accumulation of autophagosomes in the synaptic regions (from an average of 20.2 autophagosomes in *sand-1;unc-16* double mutants (n=46) compared to 8.1 autophagosomes in *atg-4.2(ola316);unc-16* double mutants (n=58) and 4.7 autophagosomes in *unc-16* single mutants (n=73); Figures 6D-6F and 6H). Importantly, early autophagy mutants like *atg-2* (required for autophagosome closure) and *atg-4.1* (required for LGG-1 priming) fail to enhance the neurite accumulation phenotype observed in *unc-16* mutant animals (Figure 6H). Together our findings indicate that *atg-4.2,* like *epg-5* and *sand-1*, acts in a late stage of autophagy prior to degradation.

We then tested the acidification state of the accumulated autophagosomes in *atg-4.2* mutants by using the tandem LGG-1 marker. We observed that in *atg-4.2* mutants, 83% of autophagosomes retained GFP signal (n=469 autophagosomes) compared to 48% in wild type neurons (n=266 autophagosomes), consistent with a defect in autophagosome maturation or acidification (Figure 6I-6O). Previous studies in yeast have observed Atg8/LGG-1 mislocalization onto the vacuole and other organelles in delipidation-defective Atg4 mutants (Nakatogawa et al., 2012; Yu et al., 2012). To test if the accumulated LGG-1 puncta in *atg-4.2* mutants are acidification-defective lysosomes, we examined lysosomes in *atg-4.2* mutants. We did not observe an accumulation of lysosomes in *atg-4.2* mutants (Figure S4B), consistent with the accumulated structures representing immature autophagosomes. Together, our findings suggest that *atg-4.2* mutants are defective in acidification, resulting in an accumulation of immature autophagosomes and a failure in degradation.

Our findings suggest that the ATG-4 proteases, while partially redundant, are also uniquely required for specific steps of autophagy, with loss of either gene resulting in distinct cell biological phenotypes of autophagy in neurons. Importantly our data uncover a novel role for ATG-4.2, and indicate that ATG-4.2 cooperates with retrograde transport mechanisms to promote autophagosome acidification and degradation in the cell soma.

## DISCUSSION

Distinct steps of the autophagy pathway occur in different subcellular compartments. Previous studies demonstrated that autophagosome biogenesis in primary neurons occurs at the distal axon and follows a compartmentalized, ordered and spatially regulated process (Maday and Holzbaur, 2014). *In vivo*, autophagosome biogenesis occurs at presynaptic sites (Soukup et al., 2016; Stavoe et al., 2016) and the number of autophagosomes in the neurite correlates with the state of activity in the neuron (Figure S1 and (Shehata et al., 2012; Soukup et al., 2016; Wang et al., 2015). Autophagosomes then undergo retrograde trafficking prior to degradation (Maday et al., 2012), indicating a regional decoupling of biogenesis and degradation. In this study we extend these cell biological findings and demonstrate, using genetics, a requirement for distinct steps of autophagy in different subcellular compartments. We observe that defects in *atg-2* and *epg-6/WIP3/4*, which are required for autophagosome closure, result in accumulated autophagosomes at the synapse, while defects in *epg-5*, required for autophagosome maturation, result in accumulated autophagosomes in the cell body. Our findings extend our understanding of the spatial organization of autophagy, and underscore the importance of coordination across cellular regions and in the context of the polarized structure of the neuron.

UNC-16/JIP3-dependent retrograde transport links autophagosome biogenesis at the synapse with autophagosome degradation in the cellular soma. In *C. elegans, Drosophila* and vertebrate neurons, axonal autophagosomes are transported towards the cell body via the dynein complex (Cheng et al., 2015; Ikenaka et al., 2013; Katsumata et al., 2010; Neisch et al., 2017). UNC-16/JIP3/Sunday Driver, a motor adaptor protein that interacts with dynein, plays important and conserved roles in the regulated localization and transport of late endosomes and lysosomes, as well as other organelles (Brown et al., 2009; Byrd et al., 2001; Drerup and Nechiporuk, 2013; Edwards et al., 2015; Edwards et al., 2013; Gowrishankar et al., 2017). Our findings extend the roles for UNC-16/JIP3 and now demonstrate that it is also required for autophagosome retrograde transport. In vertebrate neurons, autophagosome retrograde transport is promoted by JIP1, a motor adaptor protein that belongs to the same family as UNC-16/JIP3 (Fu and Holzbaur, 2014; Fu et al., 2014). While we did not detect a phenotype for autophagosome retrograde transport in *C. elegans jip-1* mutants, we note that JIP1 and JIP3 share sequence and functional similarities, and are also known to cooperate in the transport of cargo (Hammond et al., 2008; Sun et al., 2017). Together, our findings ascribe an important and potentially conserved role for UNC-16/JIP3 in retrograde transport of autophagosomes in *C. elegans* neurons.

Impaired retrograde transport in *unc-16/jip3* mutants results in accumulation of LGG-1 containing structures in the neurites. In vertebrate neurons, disruption of retrograde transport also causes accumulation of autophagosomes and lysosomes in swollen neuronal processes and synapses. This, in turn, results in autophagic stress and in neurodegeneration phenotypes similar to those seen in the neurons of Alzheimer’s disease patients (Ikenaka et al., 2013; Lee et al., 2011; Nixon et al., 2005; Takats et al., 2013). Specific disruption of JIP3 in vertebrate neurons has also been associated with an increase in soluble Aβ levels, plaque size, plaque abundance, and axonal dystrophy (Gowrishankar et al., 2017). Our findings in *C. elegans* neurons are consistent with these studies and underscore the importance of retrograde transport in linking the mechanisms of autophagosome biogenesis at the synapse with degradation in the cell body. Furthermore, our observations that combined defects in both retrograde transport and degradation machinery can enhance autophagosome accumulation in the neurites raises the question of whether the accumulated autophagosomes seen in progressive neurodegenerative diseases might result from an accruement of distinct dysfunctions in different stages of autophagy over time.

ATG-4.2 genetically cooperates with UNC-16/JIP3 in the clearance of autophagosomes from axons. Our forward genetic screens revealed that lesions in *atg-4.2* enhance the autophagosome accumulation phenotype observed for *unc-16/jip3* mutants (Figure 7B-7E). ATG-4.2 is one of two Atg4 cysteine proteases in *C. elegans* known to cleave LGG-1/Atg8 in autophagy (Wu et al., 2012). We find that the other Atg4 cysteine protease in the nematode, ATG-4.1, does not enhance the autophagosome accumulation phenotype observed for *unc-16/jip3* mutants, suggesting a distinct role for ATG-4.2 in the clearance of autophagosomes from axons. Atg4 cysteine proteases are required during both early and late stages of autophagy to cleave LGG-1/Atg8, but to achieve two distinct functions: the first cleavage (termed priming) is required for the conjugation of LGG-1/Atg8 to the autophagosomal membrane to promote autophagosome elongation, while the second cleavage (termed delipidation) occurs prior to degradation, when LGG-1/Atg8 is removed from the autophagosome membrane (Kirisako et al., 2000; Maruyama and Noda, 2017). Our *in vivo* findings suggest that these distinct cleavage roles for Atg4 in LGG-1/Atg8 priming and delipidation might be regulated by distinct Atg4 proteases, and that the *C. elegans* ATG-4.2 has a specialized role in the delipidation of LGG-1/Atg8 from autophagosomal membranes. Consistent with this idea, a recent biochemical study examining mammalian Atg4 genes revealed that different Atg4s possess different activities for cleaving soluble or membrane-bound GABARAP/Atg8/LGG-1 (Kauffman et al., 2018). Moreover, biochemical characterizations of *C. elegans atg-4.1* and *atg-4.2* gene functions demonstrated that *atg-4.2* mutants accumulate more lipidated LGG-1 compared to wild type or *atg-4.1* mutant animals, consistent with distinct biochemical properties between ATG-4.1 and ATG-4.2, and with a preferential role for ATG-4.2 in delipidation (Wu et al., 2012). We now show that the different biochemical activities of ATG-4.1 and ATG-4.2 confer different requirements *in vivo*, resulting in distinct phenotypes regarding the accumulation of autophagosomes in neurons. The distinct ATG4 genes therefore differentially interact with *unc-16/jip3* mutants in enhancing defects associated with clearance of autophagosomes from neurites.

**Figure 7.**
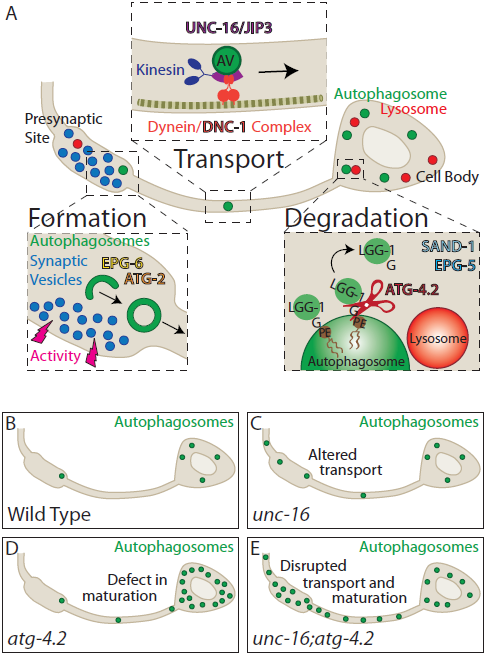
Synaptic autophagy: a model from birth at the synapse to breakdown in the cell body. (A) Model schematic illustrating the steps of synaptic autophagy. <u>Formation</u>: Autophagosomes form in the presynaptic site, influenced by the state of synaptic activity of the neuron (Figure S1 and (Shehata et al., 2012; Soukup et al., 2016; Wang et al., 2015)). Previous studies have shown that autophagy machinery is recruited in an ordered assembly to developing autophagosomes in axon tips (Maday and Holzbaur, 2014) and that the transport of transmembrane protein ATG-9, which depends on the kinesin KIF1A/UNC-104, is required to support local autophagosome biogenesis (Stavoe et al., 2016). Here we find that autophagy components ATG-2 and EPG-6/WIPI3/4, which regulate autophagosome closure, are required to initiate retrograde transport. Our LGG-1 tandem-marker indicates that synaptic autophagosomes are immature. While our data does not exclude the possibility of fusion with late endosomes or lysosomes at the synapse as previously reported (Cheng et al., 2015; Maday et al., 2012), it does indicate that autophagosome maturation is not required for retrograde transport, and that it occurs primarily during or after transport into the cell body. <u>Transport</u>: Autophagosomes undergo retrograde transport towards the cell body, a process dependent on the dynein activator dynactin/DNC-1, and the motor adaptor UNC-16/JIP3. <u>Degradation</u>: Once in the cell body, the autophagy protease ATG-4.2 cleaves LGG-1 off the autophagosome membrane to facilitate maturation and degradation in a process that also requires the RAB-7 activator SAND-1/Mon1 and effector EPG-5. (B-E) Schematics of autophagosome localization in wild type (B), *unc-16* mutant (C), *atg-4.2* mutant (D), and *unc-16;atg-4.2* double mutant (E) animals. Briefly, our data demonstrate that disruption of retrograde transport (*unc-16* mutant) or maturation (*atg-4.2* mutant) leads to autophagosome accumulation in the neurite or the cell body respectively, and that these pathways cooperate to remove autophagosomes from the synapse (E).

ATG-4.2 is necessary for autophagosome maturation and degradation. In *atg-4.2* mutants we observe the accumulation of non-acidic LGG-1 puncta in the cell bodies of neurons. These data further indicate a role for ATG-4.2 in the delipidation of LGG-1, and suggest that delipidation is necessary for the proper fusion of autophagosomes with lysosomes, and degradation. Our findings extend studies in yeast, where it has been hypothesized that delipidation of Atg8/LGG-1 from autophagosomes is necessary both for autophagosome biogenesis (Nair et al., 2012; Nakatogawa et al., 2012) and to promote fusion with lysosomes and degradation (Yu et al., 2012). Our findings further suggest that association of LGG-1 with the autophagosomal membrane, a process mediated by proteases ATG-4.1 and ATG-4.2, “bookends” the progression of autophagy. Our findings are consistent with a model (presented in Figure 7A) in which, upon biogenesis at the synapse, LGG-1 becomes conjugated to autophagosomes, a process that requires priming activity, likely performed by the priming protease ATG-4.1 to allow lipidation at the synapse. After transport to the cell body, ATG-4.2 then preferentially mediates removal of LGG-1 from the autophagosome membrane, a step that is necessary for maturation and degradation in the cell body. Therefore the specialized regulation of these proteases in distinct subcellular compartments contributes to the coordination of the autophagy pathway across the subcellular regions of the neuron.

## EXPERIMENTAL PROCEDURES

### Strains and genetics

All *C. elegans* strains were raised on NGM plates with OP50 *Escherichia coli* at 20°C, unless otherwise noted. We used N2 Bristol as the wild type reference strain. We obtained the following mutant strains through the *Caenorhabditis* Genetics Center (CGC): *dnc-1(or404)*, *unc-16(ju146), unc-16(n730), unc-16(e109), jip-1(gk466982), unc-116(e2310), unc-14(e57), atg-4.1(gk127286), atg-4.1(bp501), atg-4.2(gk430078)*, atg-4.2(gk628327), *sand-1(or552),* and *unc-13(e450).* We also obtained from the CGC, the balancer nT1[qIs51] isolated from VC3476 and used to assess escapers from *olaIs35;atg-4.1;atg-4.2/nT1*, which is sterile in its unbalanced state. We obtained mutants from Dr. Hong Zhang at the Institute of Biophysics, Chinese Academy of Sciences: *atg-2(bp576)* and *epg-6(bp242)*. And we obtained from the Mitani laboratory at the Tokyo Women’s Medical University School of Medicine: *epg-5(tm3425)*.

We also utilized *atg-9(wy56)*, DCR4750 (*olaIs35* [Pttx-3::egfp::lgg-1; Pttx-3::mCh]), and TV392 (*wyIs45* [Pttx-3:gfp:rab-3]), as described in previous publications (Colon-Ramos et al., 2007; Stavoe et al., 2016).

### Molecular biology and transgenic lines

The plasmids used in this study are derived from the pSM vector (Shen and Bargmann, 2003). We created transgenic strains using standard injection techniques. We used the following plasmids as co-injection markers: *Punc-122::gfp* (10-15ng/ul) and *Punc-122::dsRed* (30ng/ul), and generated these arrays: *olaEx3986* [Pttx-3::mCh::rab-3 (30ng/ul)], *olaEx3331* [Pttx-3::laat-1::gfp (1ng/uL), pttx-3::mCh (30ng/ul)], *olaEx3013* [Pttx-3::mCh::egfp::lgg-1) (1ng/ul)], *olaEx3015* [Pttx-3::mCh::egfp::lgg-1(G116A)) (1ng/ul)], *olaEx3430* and *olaEx3438* [pttx-3::atg-4.2 (30ng/ul)], *olaEx3426* [Pttx-3::atg-4.2(G1117A) (30ng/ul)], *olaEx3465* [Pmod-1::HisCl (30ng/ul)], *olaEx3358* [Pttx-3::ebp-2::gfp (5ng/ul), pttx-3::mCh (30ng/ul)], *olaEx1706* [Pttx-3::SP12::gfp (5ng/ul)], *olaEx1700* [Pttx-3:aman-2:gfp (0.5 ng/ul)], *olaEx1951* [Pttx-3::tom-20::gfp (30 ng/uL), pttx-3:mCh:rab-3 (30 ng/uL)], *olaIs44* [Pttx-3::egfp::lgg-1 (15ng/ul)], *olaIs58* [Paex-3::egfp::lgg-1 (15ng/ul)].

We also generated *olaEx3293* [Pelt-7::gfp (10ng/ul)] to distinguish heterozygous cross offspring from self progeny for the dominant/recessive tests.

For the cDNA constructs generated for use in this study (ATG-4.2, LAAT-1 and EBP-2), we PCR amplified cDNA from a mixed stage population of *C. elegans*. A Q5 mutagenesis kit (NEB) was then used to introduce the DNA mutation G1117A (which corresponds to the allelic lesion in *atg-4.2(ola316)*) into the ATG-4.2 cDNA. The introduction of this lesion should result in expression of the mutant protein ATG-4.2(G373R). Detailed sub-cloning information is available upon request.

### Fluorescence microscopy and confocal imaging

We used an UltraView VoX spinning disc confocal microscope with a 60x CFI Plan Apo VC, NA 1.4, oil objective on a NikonTi-E stand (PerkinElmer) with a Hammamatsu C9100-50 camera. We imaged the following fluorescently tagged fusion proteins, eGFP, GFP, RFP, and mCherry at 488 or 561 nm excitation wavelength. We anesthetized *C. elegans* at room temperature in 10mM levamisole (Sigma) or as indicated.

Images were obtained using Volocity software (Improvision by Perkin Elmer) and processed using Adobe Photoshop CS4 and (Fiji is Just) ImageJ (FIJI) software. Image processing included maximal projection, rotation, cropping, brightness/contrast, pseudo coloring, and making surface plots. Kymographs were made using the “reslice” function in FIJI. Between genotypes for compared groups the confocal laser and camera settings and the FIJI brightness/contrast settings were kept identical. All quantifications from confocal images were conducted on maximal projections of the raw data. All images are oriented anterior to the left and dorsal up.

### SNP Mapping and Whole-Genome Sequencing

We obtained mutant allele *atg-4.2(ola316)* from a visual forward genetic enhancer screen in the *unc-16(ju146);olaIs35* mutant background (integrated line *olaIs35* contains Pttx-3::egfp::lgg-1; Pttx-3::mCh). Ethyl methanesulfonate (EMS) mutagenesis was performed and animals were screened for enhanced number of autophagosomes (visualized with GFP::LGG-1) in the AIY neurite. F2 progeny were viewed on a Leica DM 5000 B compound microscope with an HCX PL APO 63x/1.40-0.60 oil objective.

The novel lesion *ola316* was out-crossed from the *unc-16* lesion and found to have an independent dominant cell body accumulation of autophagosomes. We then used single-nucleotide polymorphism (SNP) mapping as described (Davis and Hammarlund, 2006; Davis et al., 2005) to map the *ola316* lesion. Briefly, this involves crossing mutants to a divergent strain, Hawaiian CB4865, and detecting sites of recombination via comparison of dissimilar SNPs. To avoid selecting heterozygous recombinants of the dominant lesion, we mapped loss of the *ola316* locus by selecting animals with the wild type phenotype. SNP mapping was then repeated for verification by selecting and confirming homozygous animals with the *unc-16;ola316* enhanced phenotype.

We then performed whole-genome sequencing on both *unc-16;ola316;olaIs35* and for comparison *unc-16;olaIs35* animals at the Yale Center for Genome Analysis (YCGA), as previous (Sarin et al., 2008). We analyzed the results with www.usegalaxy.org, the “Cloudmap Unmapped Mutant workflow (w/ subtraction of other strains),” (Minevich et al., 2012) and verified lesions by Sanger sequencing.

### Quantification of autophagosomes in AIY

Quantifications were performed taking into consideration best practices as suggested in (Landis et al., 2012), including randomization, blinding, and data handling procedures. Particulars for each assay as follows:

#### Tandem marker

Autophagosome maturation was assessed using extrachromosomal line *olaEx3013* (Pttx-3::mCh::egfp::lgg-1).The tandem marker used in this study is based on similar probes used in mammalian neurons (Kimura et al., 2007) and in *C. elegans* tissues (Chang et al., 2017). Quantifications of puncta were performed on maximal projections of confocal micrographs and imaging settings were kept identical between compared groups. Puncta were scored as “Dual/GFP” (immature) if green puncta could be detected and as “mCh only” (mature) if the puncta displayed primarily red signal.

To ensure that the tandem labeled LGG-1 probe is lipidated onto membranes and to control for the possibility of marker aggregation, we also examined a mutant version of the probe *olaEx3015* (Pttx-3::mCh::egfp::lgg-1(G116A)), which prevents LGG-1 from associating with the autophagosomal membrane (Mizushima et al., 2010; Zhang et al., 2015). As expected, we observed that in the LGG-1(G116A) mutant probe GFP was diffuse throughout the neuron (data not shown). We also observed mCherry puncta in the cell body (3.7 puncta on average, n=34 neurons), consistent with an accumulation of mCherry (which is stable at low pH (Shaner et al., 2004)), in a lysosomal compartment. Importantly our observations suggest that the tandem marker is lipidated at the G116 residue and is expected to associate with autophagosomes as described (Mizushima et al., 2010; Zhang et al., 2015).

#### Autophagosome dynamics

To assess autophagosome dynamics over time, we used the integrated line *olaIs35* (Pttx-3::egfp::lgg-1; Pttx-3::mCh). From a dataset described previously (Stavoe et al., 2016), where confocal z-stacks were acquired once per minute for 30 minutes in wild type and autophagy mutant animals, we performed a new analysis, tracking individual autophagosomes over time and scoring the percentage of autophagosomes retained in the presynaptic region for the full 30 minutes. We also collected a new dataset, imaging at maximal speed for 5 minutes in wild type (n=71 neurons) and *unc-16(ju146)* mutant animals (n=73 neurons). These videos were scored for percentage of autophagosomes present in each of the three sub-neurite zones (Colon-Ramos et al., 2007), for anterograde and retrograde directionality of autophagosome trafficking in the neurite, and for percentage of autophagosomes leaving the cell body towards the neurite, and then returning to the cell body within the imaging window of 5 minutes.

#### Autophagosome accumulation in mutant neurites and cell bodies

To investigate the accumulation of autophagosomes in the AIY neuron, we tested mutant alleles in the *olaIs35* background. For the temperature sensitive *dnc-1(or404)* allele (Koushika et al., 2004) and a wild type control, animals were held at either 20C (permissive) or 25C (restrictive) for 48 hours prior to examination. Other candidate transport mutants (*unc-16*, *jip-1*, *unc-116*, *unc-14* and controls) were kept at 20C. Autophagosome accumulation in the neurite was then scored on a Leica DM 5000 B compound microscope. For each animal, the sum of autophagosomes in both AIY neurons was counted, then divided by two and reported as an average per neuron in Figure 3. Additional quantifications (presented in figures 4, 5, and 6) for the presence of autophagosomes in the cell bodies and neurites of mutant animals (*unc-16*, *atg-4.2*, *atg-4.1*, *epg-5*, *sand-1* etc.), double mutant animals, and animals overexpressing ATG-4.2 arrays, were scored as follows: raw z-stacks were divided to separate individual AIY neurons, maximal projections were created, and images were randomized and scored blindly.

#### Activity-dependent autophagy

To assess synaptic autophagy, animals containing *olaIs35* in a wild type, *atg-9*, *atg-2*, or *unc-13* mutant background were grown at 20C and then shifted to 25C for variable lengths of time (as indicated) and assessed for number of autophagosomes in the AIY neurite on a Leica DM 5000 B compound microscope. For the HisCl experiments, the animals additionally contained the extrachromosomal array *olaEx3465* (Pmod-1::HisCl) and were transferred to 10mM Histamine-containing or control NGM plates at 20C or 25C for 4 hours prior to assessment (Pokala et al., 2014). Animals were then mounted in either 5mM levamisole in M9 buffer (compared to 10mM used in all other experiments) or in M9 buffer containing 5mM levamisole and 10mM Histamine and scored blindly. Results are reported in Figure S1.

### Other Quantifications in AIY

#### Lysosomes

To quantify lysosome number in AIY, we observed the extrachromosomal transgenic line *olaEx3331* (Pttx-3::laat-1::gfp, pttx-3::mCh) in wild type or mutant backgrounds on a Leica DM 5000 B compound microscope. The number of GFP puncta in the two AIY neurons in the animal were counted by location, divided by two and then reported as an average per neuron.

#### Endoplasmic reticulum (ER), Golgi, and Mitochondria

For examination of ER, Golgi, and mitochondria the following extrachromosomal lines were used, respectively: *olaEx1706* (Pttx-3::SP12::gfp), *olaEx1700* (Pttx-3:aman-2:gfp), and *olaEx1951* (Pttx-3::tom-20::gfp, pttx-3:mCh:rab-3) (Abe et al., 2000; Qi et al., 2012; Rolls et al., 2002) in wild type and *unc-16(ju146)* mutant backgrounds. Quantifications were performed on maximal projection confocal micrographs. Fluorescence intensity was measured by tracing the neurite and background in FIJI (Schindelin et al., 2012) and reported as an average of background-subtracted fluorescence intensity. The percent of the neurite occupied by mitochondria was calculated as a ratio of length of the neurite with background-subtracted fluorescence signal greater than zero divided by the total length of the neurite.

#### Presynaptic enrichment

To quantify presynaptic enrichment in AIY, we used the integrated transgenic line *wyIs45,* expressing pttx-3::gfp::rab-3 in wild type and mutant backgrounds, as described (Colon-Ramos et al., 2007; Stavoe et al., 2016). The AIY neurite is divided into three zones as follows, consistent with the *C. elegans* EM reconstruction (White et al., 1986): Zone 1 is proximal to the cell body and asynaptic; Zone 2 is defined morphologically as a ∼5-μm synapse-rich region at the dorsal turn of the neuron into the nerve ring; Zone 3 is the distal part of the neurite that traverses the nerve ring and contains punctate synaptic sites. We calculated the GFP::RAB-3 enrichment using maximal projection confocal micrographs and measured fluorescence intensity across presynaptic Zones 2 and 3 with the line scan function in FIJI (Schindelin et al., 2012). Fluorescence intensity was background subtracted. All settings, for the confocal microscope and camera, were kept identical between genotypes, and neurons were only quantified when all neurite fluorescence signal was within the measurable range. Synaptic enrichment was reported as signal in Zone 2 (the first 20% of the synaptic region) divided over Zones 2 and 3 (as described in (Stavoe and Colon-Ramos, 2012)) and reported as a percentage and shown in Figure S6.

#### Microtubule EBP-2 quantification

To examine microtubule polarity, we analyzed the direction of movement for EBP-2 comets (Baas and Lin, 2011; Maniar et al., 2011), which track with the growing ends of microtubules, along the AIY neurite using the extrachromosomal line *olaEx3358* (Pttx-3::ebp-2::gfp, pttx-3::mCh). Confocal z-stacks were acquired at maximal speed for one minute and comets were assessed for movement towards the cell body (minus end out microtubule) or movement away from the cell body (plus end out microtubule) in wild type (n=18 animals) and *unc-16* mutants (n=16 animals).

### Statistical Analyses

Statistical analyses were conducted with PRISM software. For each case, the chosen statistical test is described in the figure legend. Briefly, for continuous data and for cases of counting structures, comparisons between two groups were determined by the Student’s t test, while groups of three or more were analyzed with a one-way ANOVA with *post hoc* analysis by Turkey’s multiple comparison test. Error bars were reported as standard errors of the mean (SEM). For categorical data, groups were compared with Fisher’s exact test, or for samples corresponding to large datasets, the chi-square test was used. Errors show a 95% confidence interval.

## AUTHOR CONTRIBUTIONS

SEH and DACR designed the experiments; SEH performed the experiments and data analyses. SEH and DACR prepared the manuscript.

## ACKNOWLEDGEMENTS

We thank members of the Colón-Ramos lab, the Thomas Melia lab (Yale University), Shawn Ferguson (Yale University) and Andrea Stavoe (University of Pennsylvania) for their thoughtful comments on the project. We thank undergraduate researchers Alec Rodriguez (Yale University) and Enrique Cruz-Reyes (Universidad de Puerto Rico, Cayey) and Sisi Yang (Colón-Ramos lab) for experimental assistance. We thank the *Caenorhabditis* Genetics Center (supported by the National Institutes of Health (NIH), the Office of Research Infrastructure Programs; P40 OD010440) for strains, the Mitani laboratory of the Tokyo Women’s Medical University School of Medicine for autophagy mutant strains and the Hong Zhang laboratory at the Institute of Biophysics, Chinese Academy of Sciences for autophagy mutant alleles. We thank Cori Bargmann for the HisCl construct (Pokala et al., 2014). We thank Z. Altun (www.wormatlas.org) for diagrams used in figures. We thank the Research Center for Minority Institutions program and the Instituto de Neurobiología de la Universidad de Puerto Rico for providing a meeting and brainstorming platform. This work was partially conducted at the Marine Biological Laboratories at Woods Hole under a Whitman research award to D.A.C.-R. Support for S.E.H. was provided by the Cellular and Molecular Biology Training grant T32-GM007223 from the NIH and the NSF Graduate Research Fellowship DGE-1122492. Research was supported by a Howard Hughes Medical Institute Scholar Award, by a National Science Foundation grant (NSF IOS 1353845) and the National Institutes of Health (R01NS076558).

## Author Information

The authors declare no competing financial interests. Correspondence and request for materials should be addressed to D.A.C-.R.

## SUPPLEMENTAL FIGURE LEGENDS

**Supplemental Figure 1.**
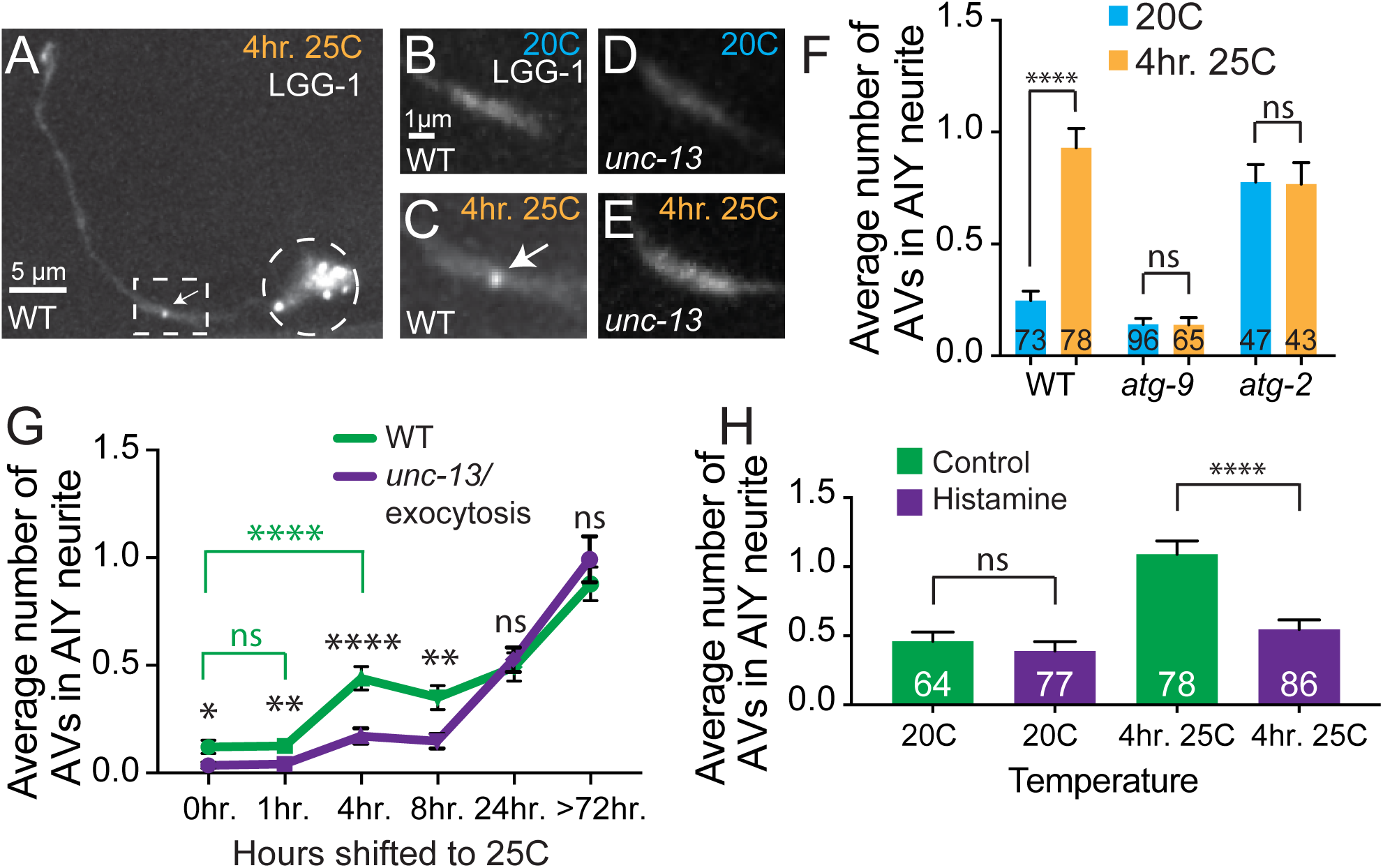
Synaptic autophagy requires neuronal activity. (A-E) Autophagosomes (visualized with GFP::LGG-1) in a representative wild type AIY neuron held at 25C for 4 hours prior to imaging, or at 20C (control). The AIY interneurons are part of the thermotaxis circuit in *C. elegans*, and the activity states of the AIYs are known to increase when animals are held at 25C (as compare to animals at 20C) (Biron et al., 2006; Clark et al., 2006; Hawk et al., 2018). We therefore used the physiological stimuli of temperature in this intact circuit to increase the activity state of the AIY neuron and examine synaptic autophagy *in vivo* under these conditions. In B-E, the synaptic-rich region of AIY in wild type (B and C) and *unc-13(e450)* mutant (D and E) neurons are shown in animals held at 20C (B and D) or held at 25C for 4 hours prior to imaging (C and E). The synaptic region of (A) (dashed boxed) is enlarged in (C). A dashed circle encloses the cell body in (A). Arrows denote autophagosomes in the synaptic region. Abbreviation: “hr.” is hours. Scale bar in (A) for (A); in (B) for (B)-(E). (F) Quantification of the average number of autophagosomes in the neurite of AIY in wild type and two autophagy mutants, *atg-9(wy56)* and *atg-2(bp576)*, held at 20C (blue bars) or 25C for 4 hours (yellow bars) prior to quantification. Note how 4-hour exposure to 25C, which increases the activity state of the AIY interneuron (Biron et al., 2006; Hawk et al., 2018), results in a higher average number of autophagosomes in the neurite. This increase is dependent on components of the autophagy pathway, as mutations that disrupt autophagosome biogenesis (*atg-9* mutants), or autophagosome closure (*atg-2* mutants) suppress the increase in autophagosomes seen for animals exposed to 25C for 4 hours. The higher number of LGG-1 puncta seen in *atg-2* mutants at 20C is likely due to accumulation of unclosed autophagosomes which, as previously reported, suppresses autophagosome biogenesis (Stavoe et al., 2016). Error bars show standard error of the mean (SEM). ****p<0.0001 by one-way ANOVA with Tukey’s post hoc analysis between 20C and 25C conditions. Abbreviations: “ns” is not significant; “hr.” is hours; “AV” is autophagic vacuole. The number of animals examined is shown on each bar. (G) Quantification of the average number of autophagosomes in the AIY neurite in wild type (green line) and exocytosis/*unc-13(e450)* mutant (purple line) animals. Animals were raised at 20C and then held at 25C for variable lengths of time as indicated. Note that after 4 hours at 25C, higher numbers of synaptic autophagosomes can be observed, and that this increase is suppressed in exocytosis/*unc-13(e450)* mutant animals (purple line). Our findings also demonstrate activity independent increases in autophagosome number for animals raised at 25C for 24 hours or longer, likely due to temperature-dependent effects in the cell and consistent with the known detrimental effects of prolonged exposure of *C. elegans* to 25C temperature conditions (Byerly et al., 1976; Hosono et al., 1982). For all genotypes and time points, number of animals examined is n≥45. *p<0.05, **p<0.01, ****p<0.0001 by Student’s t test between wild type and *unc-13* mutant animals at each time point (black significance values) or between different wild type time points (green significance values). Abbreviations: “ns” is not significant between wild type and *unc-13* mutants or as indicated; “hr.” is hour(s); “AV” is autophagic vacuole. (H) Quantification of the average number of autophagosomes in the AIY neurite in animals expressing the exogenous inhibitory chloride channel (HisCl) (Pokala et al., 2014) when exposed to Histamine (purple bars) or control (green bars). Consistent with the suppression observed for exocytosis/*unc-13(e450)* mutant animals in (G), these findings demonstrate that the increase in autophagosome number observed for animals held for 4 hours at 25C can be suppressed by inhibiting synaptic activity, and suggest a link, under physiological conditions *in vivo*, between the activity state of the neuron and the number of neuronal autophagosomes. Error bars show standard error of the mean (SEM). ****p<0.0001 by one-way ANOVA with Tukey’s post hoc analysis between Histamine and control conditions. Abbreviations: “ns” is not significant; “AV” is autophagic vacuole. The number of animals examined is shown on each bar.

**Supplemental Figure 2.**
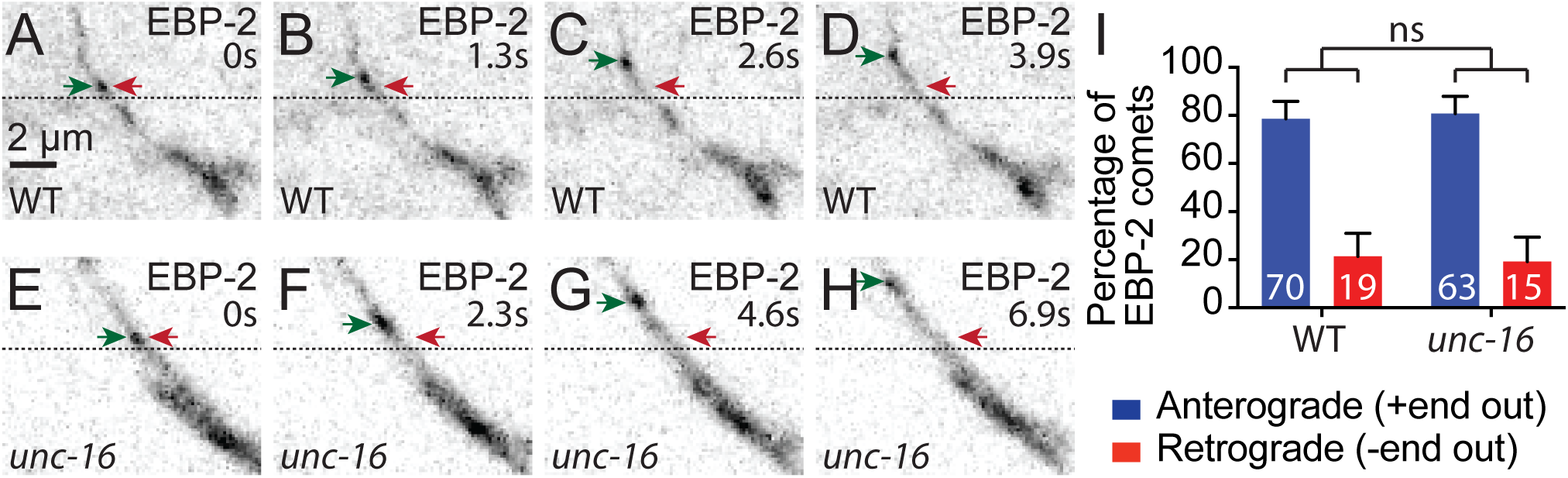
Microtubule polarity in AIY is unaltered in *unc-16/jip3* mutants. (A-H) Time series of representative growing (+) ends of microtubules, visualized with (EBP-2::GFP) in wild type (A-D) and *unc-16(ju146)/jip3* (E-H) mutant animals. We include dashed lines at the same position and across time points for reference. Red arrows denote EBP-2 puncta start positions. Green arrows track EBP-2 puncta positions over time. Time is indicated in the upper right corner of the image and “s” is seconds. Images are oriented distal axon towards the upper left, cell body towards the lower right. Scale bar in (A) for (A)-(H). (I) Quantification of the percentage of EBP-2 comets trafficking in the anterograde direction away from the cell body (corresponding microtubule is plus (+) end out) or the retrograde direction towards the cell body (corresponding microtubule is minus (-) end out) in wild type and *unc-16(ju146)* mutant animals. The number of EBP-2 comets examined is shown on each bar for ≥16 animals for each genotype. Note that the AIY neurite shows a primarily plus-end-out microtubule orientation, which is a common feature of axons (Baas and Lin, 2011), and which is not altered in *unc-16(ju146)/jip3* mutant animals. “ns,” not significant by Fisher’s exact test between wild type and mutant animals. Error bars show a 95% confidence interval.

**Supplemental Figure 3.**
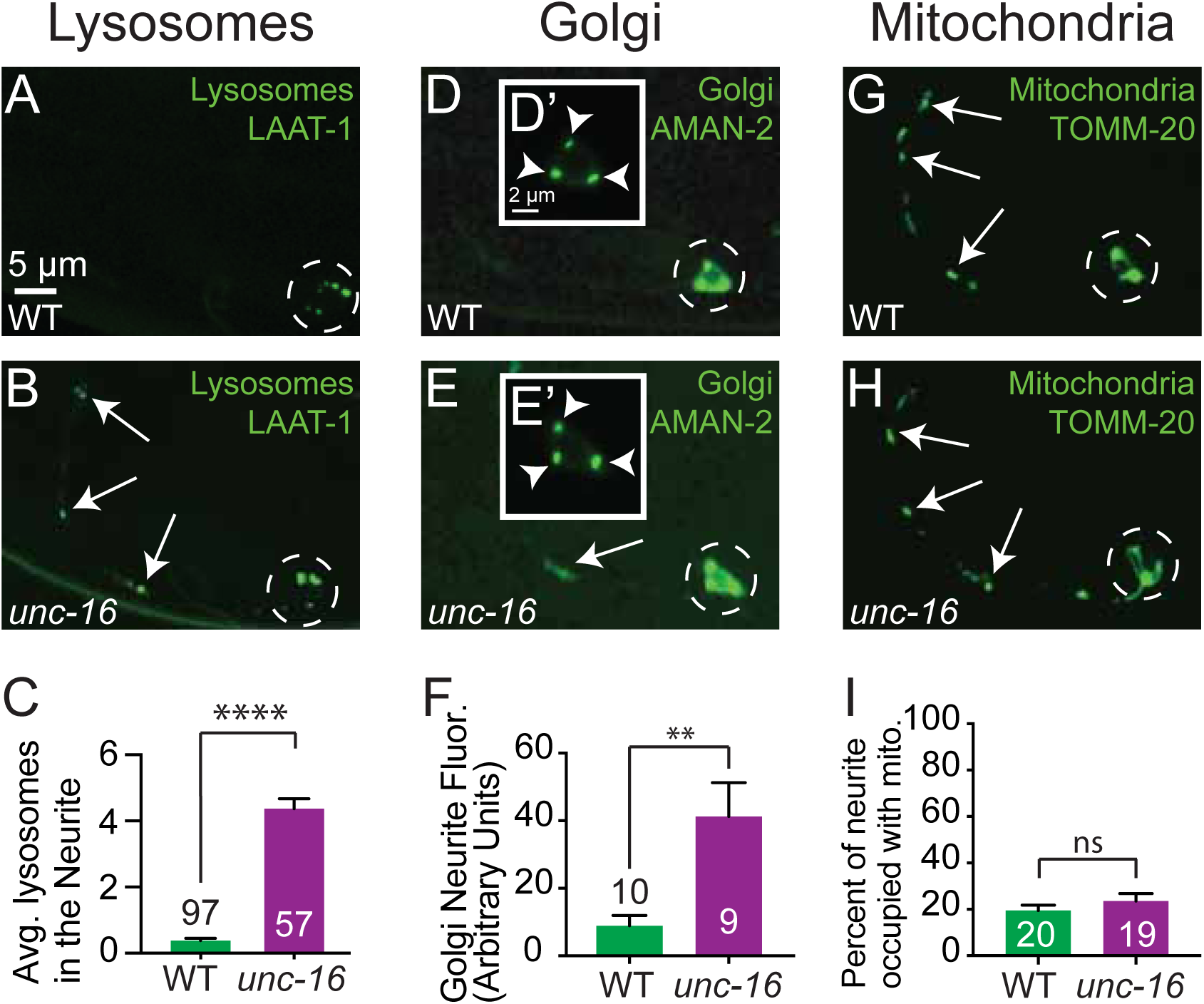
Organelle localizations in *unc-16/jip3* mutants. (A and B) Representative images of lysosomes (visualized with LAAT-1::GFP) (Liu et al., 2012) in wild type and *unc-16(ju146)* mutant animals. Dashed circles enclose the cell bodies. Arrows denote lysosomes in the neurite. (C) Quantification of the average number of lysosomes in the neurite of AIY in wild type and *unc-16(ju146)* mutant animals. Error bars show standard error of the mean (SEM). ****p<0.0001 by Student’s t test between wild type and mutants. The number of animals examined is shown on each bar. For wild type, different analyses of the same dataset are shown in figures 1P, and S4B. (D and E) Representative images of Golgi (visualized with AMAN-2::GFP) (Rolls et al., 2002) in wild type and *unc-16(ju146)* mutant animals. Dashed circles enclose the cell bodies. Arrow denotes AMAN-2::GFP in the neurite. Insets D’ in D and E’ in E show enlarged views of the cell body with different brightness and contrast as compared to the main image and adjusted to better visualize the cell body. Arrowheads denote Golgi puncta in the cell body. There was no significant difference in the number of cell body Golgi puncta between wild type (2.5 puncta, n=10 neurons) and *unc-16* (2.9 puncta, n=9 neurons) mutants (data not shown). (F) Quantification of background subtracted fluorescence signal in the neurite of AIY in wild type and *unc-16(ju146)* mutant animals. Error bars show standard error of the mean (SEM). **p<0.01 by Student’s t test between wild type and mutants. The number of neurons examined is shown on each bar. We note that the neurite signal, while brighter in *unc-16* mutants, is faint and cytoplasmic rather than the expected punctate pattern seen in the cell body for Golgi stacks. (G and H) Representative images of mitochondria (visualized with TOM20::GFP) (Abe et al., 2000; Qi et al., 2012) in wild type and *unc-16(ju146)* mutant animals. Dashed circles enclose the cell bodies. Arrows denote mitochondria in the neurite. (I) Quantification of the percentage of the length of AIY neurite occupied by mitochondria in wild type and *unc-16(ju146)* mutant animals. Neurite fluorescence intensity analyses of the mitochondria revealed higher intensity for TOM20::GFP in *unc-16* mutants, similar to the results seen for the for the Golgi AMAN-2::GFP marker, but not an increase in mitochondria occupancy in the neurite. Analyses for the endoplasmic reticulum (ER) (visualized with SP12::GFP) (Rolls et al., 2002) revealed no significant difference in ER signal intensity in the neurites of wild type and *unc-16(ju146)* mutant animals (data not shown). Error bars show standard error of the mean (SEM). “ns,” not significant by Student’s t test between wild type and mutants. The number of neurons examined is shown on each bar. Scale bar in (A) for (A), (B), (D), (E), (G), and (H); in (D’) for (D’) and (E’).

**Supplemental Figure 4.**
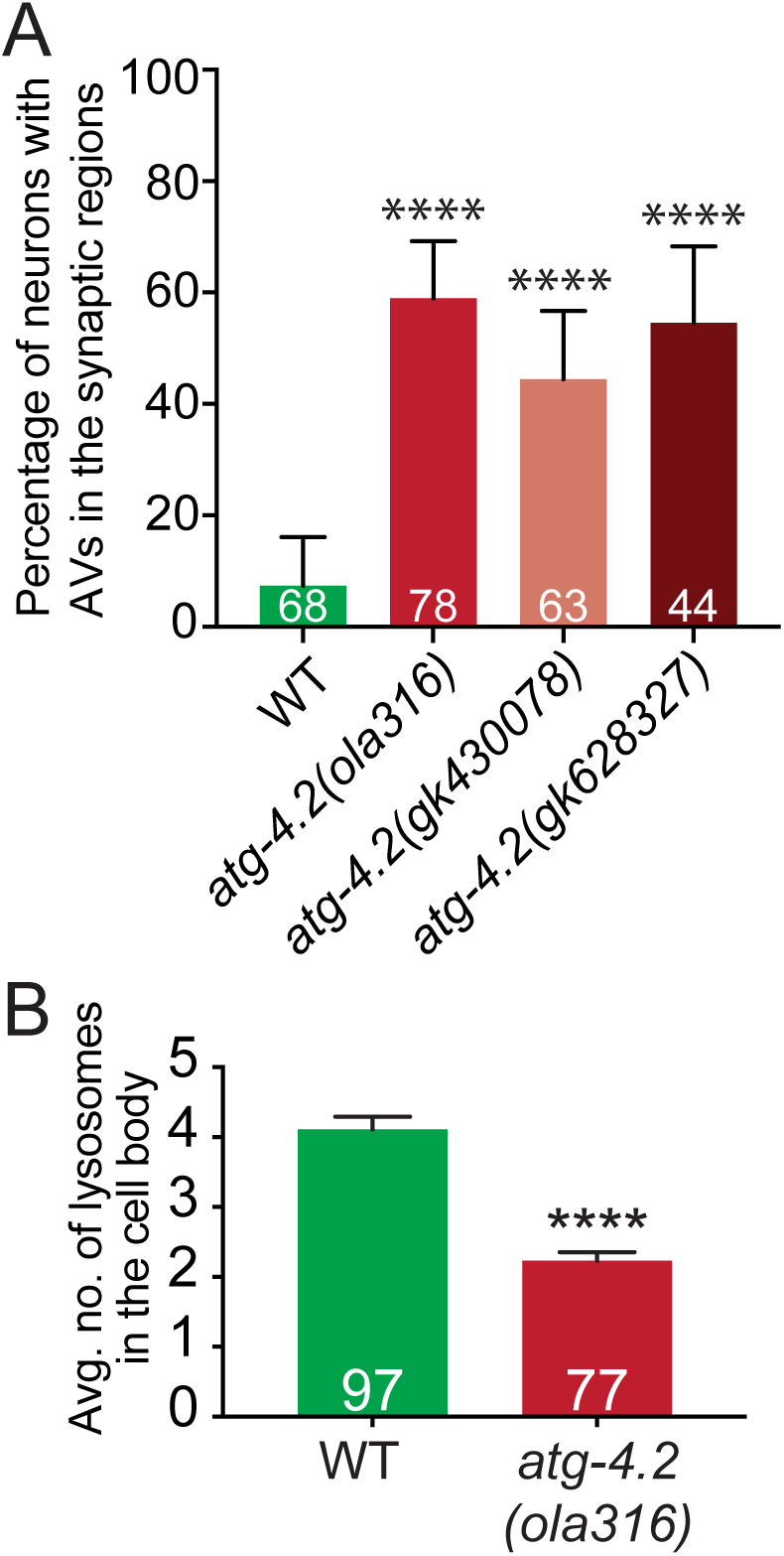
Quantification of autophagosome and lysosome phenotypes in the neurites and cell bodies of *atg-4.2* mutant animals. (A) Quantification of the percentage of neurons with autophagosomes in the synaptic regions of AIY in wild type, and three independent alleles of *atg-4.2 (ola316, gk430078,* and *gk628327)*. While all alleles of *atg-4.2* mutant animals display an increase in the number of neurons with autophagosomes in the neurite (as compared to wild type), we note that the expressivity of this phenotype (the number of autophagosomes per synaptic region) is low as compared to *unc-16* mutant animals. ****p<0.0001 by Fisher’s exact test between wild type and mutant animals. Error bars show a 95% confidence interval. The number of neurons examined is shown on each bar. (B) Quantification of the average number of lysosomes in the cell body of wild type (green bar) and *atg-4.2(ola316)* mutant (red bar) animals visualized using the LAAT-1::GFP marker (Liu et al., 2012). The observation that *atg-4.2* mutants display fewer lysosomes on average as compared to wild type could be due to a consequence of *atg-4.2* in lysosome biogenesis, turnover, or an indirect consequence of the increase in immature autophagosomes. Error bars show standard error of the mean (SEM). ****p<0.0001 by Student’s t test between wild type and mutants. The number of animals examined is shown on each bar. For wild type, different analyses of the same dataset are shown in figures 1P, and S4B.

**Supplemental Figure 5.**
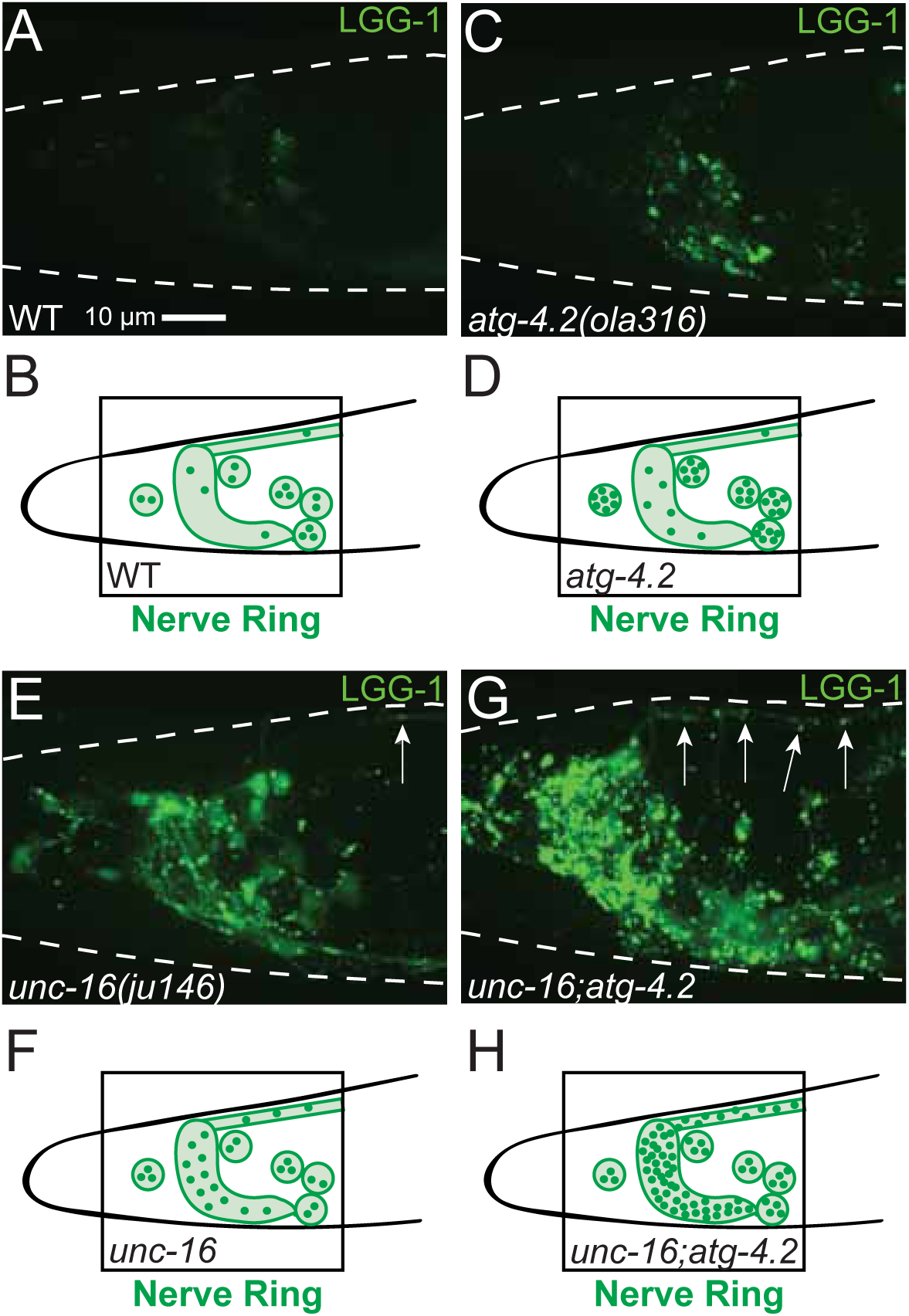
Panneuronal examination of autophagosomes in *unc-16* and *atg-4.2* mutant animals. (A-H) Panneuronal visualization of autophagosomes in the head of the worm, outlined with white dashed lines, and labeled by *olaIs58* (Paex-3::egfp::lgg-1) (A, C, E, and G). The confocal micrographs shown here in A, C, E and G include the nerve ring region and the anterior portion of the dorsal nerve cord, with schematics summarizing observations in B, D, F and H. Results shown for wild type (A and B), *atg-4.2(ola316)* (C and D), *unc-16(ju146)* (E and F), and *unc-16(ju146);atg-4.2(ola316)* double (G and H) mutant animals. Note that the phenotypes reported here are consistent with those reported using AIY-specific promoters in Figure 4, indicating that these phenotypes are panneuronal. Arrows refer to autophagosomes in the dorsal nerve cord. In B, D, F, and H, green dots represent autophagosomes within the nerve ring, cell bodies, and dorsal nerve cord. Scale bar in (A) for (A), (C), (E), and (G).

**Supplemental Figure 6.**
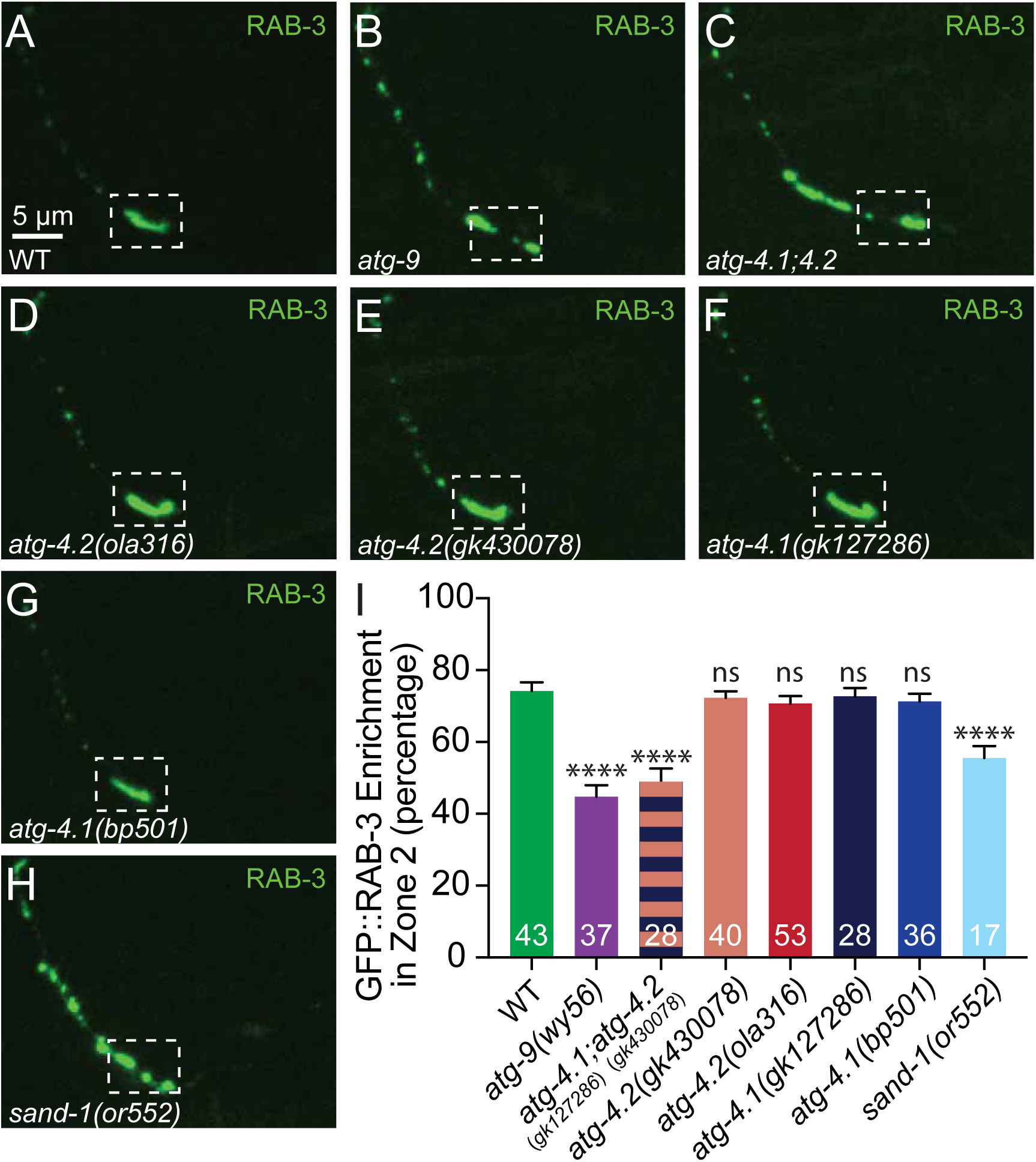
The autophagy proteases *atg-4.1* and *atg-4.2* are redundantly required for presynaptic assembly in AIY. (A-H) Representative confocal images of presynaptic structures in AIY (visualized with GFP::RAB-3 to label synaptic vesicles) in wild type (A), *atg-9(wy56)* mutant (B), *atg-4.1(gk127286);atg-4.2(gk430078)* double mutant (C), *atg-4.2(ola316)* mutant (D), *atg-4.2(gk430078)* mutant (E), *atg-4.1(gk127286)* mutant (F), *atg-4.1(bp501)* mutant (G), and *sand-1(or552)* mutant (H) animals. The dashed box encloses the synapse-enriched zone 2 region of the neurite, which displays a continuous pattern of synaptic vesicles in wild type animals. In *atg-9* mutants, *atg-4.1;atg-4.2* double mutants, and *sand-1* mutants, but not in *atg-4.1* or *atg-4.2* single mutants, note a disrupted pattern of synaptic vesicles representative of synaptogenesis defects (Stavoe et al., 2016). Scale bar in (A) for (A)-(H). (I) Quantification of synaptic enrichment in Zone 2 of the AIY neurite in wild type, *atg-9(wy56)* mutants, *atg-4.1(gk127286);atg-4.2(gk430078)* double mutants, *atg-4.2(gk430078)* mutants, *atg-4.2(ola316)* mutants, *atg-4.1(gk127286)* mutants, *atg-4.1(bp501)* mutants, and *sand-1(or552)* mutant animals. Error bars show standard error of the mean (SEM). ****p<0.0001 by one-way ANOVA with Tukey’s post hoc analysis between wild type and mutants. Abbreviations: “ns” is not significant as compared to wild type neurons. Numbers on bars represent number of neurons examined.

**Supplemental Movie S1.** Time lapse video of GFP::LGG-1 in a wild type AIY neuron showing autophagosome biogenesis and retrograde transport from the synaptic region

**Supplemental Movie S2.** Time lapse video of GFP::LGG-1 in an AIY neuron in an *atg-2(bp576)* mutant animal, showing autophagosomes in the synaptic region

**Supplemental Movie S3.** Time lapse video of GFP::LGG-1 in a pair of AIY neurons in an *epg-6(bp424)* mutant animal, showing autophagosomes in the synaptic region

**Supplemental Movie S4.** Time lapse video of GFP::LGG-1 in a pair of AIY neurons in an *epg-5(tm3425)* mutant animal, showing autophagosome retrograde transport from the synaptic region

## REFERENCES

Abe, Y., Shodai, T., Muto, T., Mihara, K., Torii, H., Nishikawa, S., Endo, T., and Kohda, D. (2000). Structural basis of presequence recognition by the mitochondrial protein import receptor Tom20. Cell 100, 551–560.

Alberti, A., Michelet, X., Djeddi, A., and Legouis, R. (2010). The autophagosomal protein LGG-2 acts synergistically with LGG-1 in dauer formation and longevity in C. elegans. Autophagy 6, 622–633.

Arimoto, M., Koushika, S.P., Choudhary, B.C., Li, C., Matsumoto, K., and Hisamoto, N. (2011). The Caenorhabditis elegans JIP3 protein UNC-16 functions as an adaptor to link kinesin-1 with cytoplasmic dynein. The Journal of neuroscience: the official journal of the Society for Neuroscience 31, 2216–2224.

Baas, P.W., and Lin, S. (2011). Hooks and comets: The story of microtubule polarity orientation in the neuron. Dev Neurobiol 71, 403–418.

Betin, V.M., Singleton, B.K., Parsons, S.F., Anstee, D.J., and Lane, J.D. (2013). Autophagy facilitates organelle clearance during differentiation of human erythroblasts: evidence for a role for ATG4 paralogs during autophagosome maturation. Autophagy 9, 881–893.

Brown, H.M., Van Epps, H.A., Goncharov, A., Grant, B.D., and Jin, Y. (2009). The JIP3 scaffold protein UNC-16 regulates RAB-5 dependent membrane trafficking at C. elegans synapses. Dev Neurobiol 69, 174–190.

Byrd, D.T., Kawasaki, M., Walcoff, M., Hisamoto, N., Matsumoto, K., and Jin, Y. (2001). UNC-16, a JNK-signaling scaffold protein, regulates vesicle transport in C. elegans. Neuron 32, 787–800.

Cavalli, V., Kujala, P., Klumperman, J., and Goldstein, L.S. (2005). Sunday Driver links axonal transport to damage signaling. The Journal of cell biology 168, 775–787.

Chang, J.T., Kumsta, C., Hellman, A.B., Adams, L.M., and Hansen, M. (2017). Spatiotemporal regulation of autophagy during Caenorhabditis elegans aging. Elife 6.

Cheng, X.T., Zhou, B., Lin, M.Y., Cai, Q., and Sheng, Z.H. (2015). Axonal autophagosomes recruit dynein for retrograde transport through fusion with late endosomes. The Journal of cell biology 209, 377–386.

Colon-Ramos, D.A., Margeta, M.A., and Shen, K. (2007). Glia promote local synaptogenesis through UNC-6 (netrin) signaling in C. elegans. Science 318, 103–106.

Davis, M.W., and Hammarlund, M. (2006). Single-nucleotide polymorphism mapping. Methods in molecular biology 351, 75–92.

Davis, M.W., Hammarlund, M., Harrach, T., Hullett, P., Olsen, S., and Jorgensen, E.M. (2005). Rapid single nucleotide polymorphism mapping in C. elegans. BMC genomics 6, 118.

Drerup, C.M., and Nechiporuk, A.V. (2013). JNK-interacting protein 3 mediates the retrograde transport of activated c-Jun N-terminal kinase and lysosomes. PLoS genetics 9, e1003303.

Edwards, S.L., Morrison, L.M., Yorks, R.M., Hoover, C.M., Boominathan, S., and Miller, K.G. (2015). UNC-16 (JIP3) Acts Through Synapse-Assembly Proteins to Inhibit the Active Transport of Cell Soma Organelles to Caenorhabditis elegans Motor Neuron Axons. Genetics 201, 117–141.

Edwards, S.L., Yu, S.C., Hoover, C.M., Phillips, B.C., Richmond, J.E., and Miller, K.G. (2013). An organelle gatekeeper function for Caenorhabditis elegans UNC-16 (JIP3) at the axon initial segment. Genetics 194, 143–161.

Fu, M.M., and Holzbaur, E.L. (2014). Integrated regulation of motor-driven organelle transport by scaffolding proteins. Trends in cell biology 24, 564–574.

Fu, M.M., Nirschl, J.J., and Holzbaur, E.L.F. (2014). LC3 binding to the scaffolding protein JIP1 regulates processive dynein-driven transport of autophagosomes. Developmental cell 29, 577–590.

Fujita, N., Hayashi-Nishino, M., Fukumoto, H., Omori, H., Yamamoto, A., Noda, T., and Yoshimori, T. (2008). An Atg4B mutant hampers the lipidation of LC3 paralogues and causes defects in autophagosome closure. Molecular biology of the cell 19, 4651–4659.

Gowrishankar, S., Wu, Y., and Ferguson, S.M. (2017). Impaired JIP3-dependent axonal lysosome transport promotes amyloid plaque pathology. The Journal of cell biology 216, 3291–3305.

Gutierrez, M.G., Munafo, D.B., Beron, W., and Colombo, M.I. (2004). Rab7 is required for the normal progression of the autophagic pathway in mammalian cells. Journal of cell science 117, 2687–2697.

Hammond, J.W., Griffin, K., Jih, G.T., Stuckey, J., and Verhey, K.J. (2008). Co-operative versus independent transport of different cargoes by Kinesin-1. Traffic 9, 725–741.

Hegedus, K., Takats, S., Boda, A., Jipa, A., Nagy, P., Varga, K., Kovacs, A.L., and Juhasz, G. (2016). The Ccz1-Mon1-Rab7 module and Rab5 control distinct steps of autophagy. Molecular biology of the cell 27, 3132–3142.

Hollenbeck, P.J. (1993). Products of endocytosis and autophagy are retrieved from axons by regulated retrograde organelle transport. The Journal of cell biology 121, 305–315.

Hyttinen, J.M., Niittykoski, M., Salminen, A., and Kaarniranta, K. (2013). Maturation of autophagosomes and endosomes: a key role for Rab7. Biochimica et biophysica acta 1833, 503–510.

Ikenaka, K., Kawai, K., Katsuno, M., Huang, Z., Jiang, Y.M., Iguchi, Y., Kobayashi, K., Kimata, T., Waza, M., Tanaka, F., et al. (2013). dnc-1/dynactin 1 knockdown disrupts transport of autophagosomes and induces motor neuron degeneration. PloS one 8, e54511.

Itakura, E., Kishi-Itakura, C., and Mizushima, N. (2012). The hairpin-type tail-anchored SNARE syntaxin 17 targets to autophagosomes for fusion with endosomes/lysosomes. Cell 151, 1256–1269.

Jager, S., Bucci, C., Tanida, I., Ueno, T., Kominami, E., Saftig, P., and Eskelinen, E.L. (2004). Role for Rab7 in maturation of late autophagic vacuoles. Journal of cell science 117, 4837–4848.

Kaasinen, S.K., Harvey, L., Reynolds, A.J., and Hendry, I.A. (2008). Autophagy generates retrogradely transported organelles: a hypothesis. Int J Dev Neurosci 26, 625–634.

Katsumata, K., Nishiyama, J., Inoue, T., Mizushima, N., Takeda, J., and Yuzaki, M. (2010). Dynein- and activity-dependent retrograde transport of autophagosomes in neuronal axons. Autophagy 6, 378–385.

Kauffman, K.J., Yu, S., Jin, J., Mugo, B., Nguyen, N., O’Brien, A., Nag, S., Lystad, A.H., and Melia, T.J. (2018). Delipidation of mammalian Atg8-family proteins by each of the four ATG4 proteases. Autophagy, 1-56.

Kimura, S., Noda, T., and Yoshimori, T. (2007). Dissection of the autophagosome maturation process by a novel reporter protein, tandem fluorescent-tagged LC3. Autophagy 3, 452–460.

Kirisako, T., Baba, M., Ishihara, N., Miyazawa, K., Ohsumi, M., Yoshimori, T., Noda, T., and Ohsumi, Y. (1999). Formation process of autophagosome is traced with Apg8/Aut7p in yeast. The Journal of cell biology 147, 435–446.

Kirisako, T., Ichimura, Y., Okada, H., Kabeya, Y., Mizushima, N., Yoshimori, T., Ohsumi, M., Takao, T., Noda, T., and Ohsumi, Y. (2000). The reversible modification regulates the membrane-binding state of Apg8/Aut7 essential for autophagy and the cytoplasm to vacuole targeting pathway. The Journal of cell biology 151, 263–276.

Koushika, S.P., Schaefer, A.M., Vincent, R., Willis, J.H., Bowerman, B., and Nonet, M.L. (2004). Mutations in Caenorhabditis elegans cytoplasmic dynein components reveal specificity of neuronal retrograde cargo. The Journal of neuroscience: the official journal of the Society for Neuroscience 24, 3907–3916.

Kumanomidou, T., Mizushima, T., Komatsu, M., Suzuki, A., Tanida, I., Sou, Y.S., Ueno, T., Kominami, E., Tanaka, K., and Yamane, T. (2006). The crystal structure of human Atg4b, a processing and de-conjugating enzyme for autophagosome-forming modifiers. J Mol Biol 355, 612–618.

Landis, S.C., Amara, S.G., Asadullah, K., Austin, C.P., Blumenstein, R., Bradley, E.W., Crystal, R.G., Darnell, R.B., Ferrante, R.J., Fillit, H., et al. (2012). A call for transparent reporting to optimize the predictive value of preclinical research. Nature 490, 187–191.

Lee, S., Sato, Y., and Nixon, R.A. (2011). Lysosomal proteolysis inhibition selectively disrupts axonal transport of degradative organelles and causes an Alzheimer’s-like axonal dystrophy. The Journal of neuroscience: the official journal of the Society for Neuroscience 31, 7817–7830.

Li, M., Hou, Y., Wang, J., Chen, X., Shao, Z.M., and Yin, X.M. (2011). Kinetics comparisons of mammalian Atg4 homologues indicate selective preferences toward diverse Atg8 substrates. The Journal of biological chemistry 286, 7327–7338.

Liang, Y., and Sigrist, S. (2017). Autophagy and proteostasis in the control of synapse aging and disease. Current opinion in neurobiology 48, 113–121.

Liu, B., Du, H., Rutkowski, R., Gartner, A., and Wang, X. (2012). LAAT-1 is the lysosomal lysine/arginine transporter that maintains amino acid homeostasis. Science 337, 351–354.

Lu, Q., Yang, P., Huang, X., Hu, W., Guo, B., Wu, F., Lin, L., Kovacs, A.L., Yu, L., and Zhang, H. (2011). The WD40 repeat PtdIns(3)P-binding protein EPG-6 regulates progression of omegasomes to autophagosomes. Developmental cell 21, 343–357.

Maday, S., and Holzbaur, E.L. (2014). Autophagosome biogenesis in primary neurons follows an ordered and spatially regulated pathway. Developmental cell 30, 71–85.

Maday, S., Wallace, K.E., and Holzbaur, E.L. (2012). Autophagosomes initiate distally and mature during transport toward the cell soma in primary neurons. The Journal of cell biology 196, 407–417.

Maniar, T.A., Kaplan, M., Wang, G.J., Shen, K., Wei, L., Shaw, J.E., Koushika, S.P., and Bargmann, C.I. (2011). UNC-33 (CRMP) and ankyrin organize microtubules and localize kinesin to polarize axon-dendrite sorting. Nature neuroscience 15, 48–56.

Manil-Segalen, M., Lefebvre, C., Jenzer, C., Trichet, M., Boulogne, C., Satiat-Jeunemaitre, B., and Legouis, R. (2014). The C. elegans LC3 acts downstream of GABARAP to degrade autophagosomes by interacting with the HOPS subunit VPS39. Developmental cell 28, 43–55.

Marino, G., Uria, J.A., Puente, X.S., Quesada, V., Bordallo, J., and Lopez-Otin, C. (2003). Human autophagins, a family of cysteine proteinases potentially implicated in cell degradation by autophagy. The Journal of biological chemistry 278, 3671–3678.

Maruyama, T., and Noda, N.N. (2017). Autophagy-regulating protease Atg4: structure, function, regulation and inhibition. J Antibiot (Tokyo).

Melendez, A., Talloczy, Z., Seaman, M., Eskelinen, E.L., Hall, D.H., and Levine, B. (2003). Autophagy genes are essential for dauer development and life-span extension in C. elegans. Science 301, 1387–1391.

Minevich, G., Park, D.S., Blankenberg, D., Poole, R.J., and Hobert, O. (2012). CloudMap: a cloud-based pipeline for analysis of mutant genome sequences. Genetics 192, 1249–1269.

Mizushima, N., Yoshimori, T., and Levine, B. (2010). Methods in mammalian autophagy research. Cell 140, 313–326.

Nah, J., Yuan, J., and Jung, Y.K. (2015). Autophagy in neurodegenerative diseases: from mechanism to therapeutic approach. Mol Cells 38, 381–389.

Nair, U., Yen, W.L., Mari, M., Cao, Y., Xie, Z., Baba, M., Reggiori, F., and Klionsky, D.J. (2012). A role for Atg8-PE deconjugation in autophagosome biogenesis. Autophagy 8, 780–793.

Nakamura, S., and Yoshimori, T. (2017). New insights into autophagosome-lysosome fusion. Journal of cell science 130, 1209–1216.

Nakatogawa, H., Ichimura, Y., and Ohsumi, Y. (2007). Atg8, a ubiquitin-like protein required for autophagosome formation, mediates membrane tethering and hemifusion. Cell 130, 165–178.

Nakatogawa, H., Ishii, J., Asai, E., and Ohsumi, Y. (2012). Atg4 recycles inappropriately lipidated Atg8 to promote autophagosome biogenesis. Autophagy 8, 177-186.

Neisch, A.L., Neufeld, T.P., and Hays, T.S. (2017). A STRIPAK complex mediates axonal transport of autophagosomes and dense core vesicles through PP2A regulation. The Journal of cell biology 216, 441–461.

Nixon, R.A. (2013). The role of autophagy in neurodegenerative disease. Nature medicine 19, 983–997.

Nixon, R.A., Wegiel, J., Kumar, A., Yu, W.H., Peterhoff, C., Cataldo, A., and Cuervo, A.M. (2005). Extensive involvement of autophagy in Alzheimer disease: an immuno-electron microscopy study. Journal of neuropathology and experimental neurology 64, 113–122.

Pokala, N., Liu, Q., Gordus, A., and Bargmann, C.I. (2014). Inducible and titratable silencing of Caenorhabditis elegans neurons in vivo with histamine-gated chloride channels. Proceedings of the National Academy of Sciences of the United States of America 111, 2770–2775.

Poteryaev, D., Fares, H., Bowerman, B., and Spang, A. (2007). Caenorhabditis elegans SAND-1 is essential for RAB-7 function in endosomal traffic. EMBO J 26, 301–312.

Qi, Y.B., Garren, E.J., Shu, X., Tsien, R.Y., and Jin, Y. (2012). Photo-inducible cell ablation in Caenorhabditis elegans using the genetically encoded singlet oxygen generating protein miniSOG. Proceedings of the National Academy of Sciences of the United States of America 109, 7499–7504.

Rolls, M.M., Hall, D.H., Victor, M., Stelzer, E.H., and Rapoport, T.A. (2002). Targeting of rough endoplasmic reticulum membrane proteins and ribosomes in invertebrate neurons. Molecular biology of the cell 13, 1778–1791.

Sarin, S., Prabhu, S., O’Meara, M.M., Pe’er, I., and Hobert, O. (2008). Caenorhabditis elegans mutant allele identification by whole-genome sequencing. Nature methods 5, 865–867.

Schindelin, J., Arganda-Carreras, I., Frise, E., Kaynig, V., Longair, M., Pietzsch, T., Preibisch, S., Rueden, C., Saalfeld, S., Schmid, B., et al. (2012). Fiji: an open-source platform for biological-image analysis. Nature methods 9, 676-682.

Shaner, N.C., Campbell, R.E., Steinbach, P.A., Giepmans, B.N., Palmer, A.E., and Tsien, R.Y. (2004). Improved monomeric red, orange and yellow fluorescent proteins derived from Discosoma sp. red fluorescent protein. Nat Biotechnol 22, 1567–1572.

Shehata, M., Matsumura, H., Okubo-Suzuki, R., Ohkawa, N., and Inokuchi, K. (2012). Neuronal stimulation induces autophagy in hippocampal neurons that is involved in AMPA receptor degradation after chemical long-term depression. The Journal of neuroscience: the official journal of the Society for Neuroscience 32, 10413–10422.

Shen, K., and Bargmann, C.I. (2003). The immunoglobulin superfamily protein SYG-1 determines the location of specific synapses in C. elegans. Cell 112, 619–630.

Son, J.H., Shim, J.H., Kim, K.H., Ha, J.Y., and Han, J.Y. (2012). Neuronal autophagy and neurodegenerative diseases. Experimental & molecular medicine 44, 89–98.

Soukup, S.F., Kuenen, S., Vanhauwaert, R., Manetsberger, J., Hernandez-Diaz, S., Swerts, J., Schoovaerts, N., Vilain, S., Gounko, N.V., Vints, K., et al. (2016). A LRRK2-Dependent EndophilinA Phosphoswitch Is Critical for Macroautophagy at Presynaptic Terminals. Neuron 92, 829–844.

Stavoe, A.K., Hill, S.E., Hall, D.H., and Colon-Ramos, D.A. (2016). KIF1A/UNC-104 Transports ATG-9 to Regulate Neurodevelopment and Autophagy at Synapses. Developmental cell 38, 171–185.

Stavoe, A.K.H., and Colon-Ramos, D.A. (2012). Netrin instructs synaptic vesicle clustering through Rac GTPase, MIG-10, and the actin cytoskeleton. Journal of Cell Biology 197, 75-88.

Sun, T., Li, Y., Li, T., Ma, H., Guo, Y., Jiang, X., Hou, M., Huang, S., and Chen, Z. (2017). JIP1 and JIP3 cooperate to mediate TrkB anterograde axonal transport by activating kinesin-1. Cell Mol Life Sci 74, 4027–4044.

Takats, S., Nagy, P., Varga, A., Pircs, K., Karpati, M., Varga, K., Kovacs, A.L., Hegedus, K., and Juhasz, G. (2013). Autophagosomal Syntaxin17-dependent lysosomal degradation maintains neuronal function in Drosophila. The Journal of cell biology 201, 531–539.

Tammineni, P., Ye, X., Feng, T., Aikal, D., and Cai, Q. (2017). Impaired retrograde transport of axonal autophagosomes contributes to autophagic stress in Alzheimer’s disease neurons. Elife 6.

Velikkakath, A.K., Nishimura, T., Oita, E., Ishihara, N., and Mizushima, N. (2012). Mammalian Atg2 proteins are essential for autophagosome formation and important for regulation of size and distribution of lipid droplets. Molecular biology of the cell 23, 896–909.

Vijayan, V., and Verstreken, P. (2017). Autophagy in the presynaptic compartment in health and disease. The Journal of cell biology 216, 1895–1906.

Wang, T., Martin, S., Papadopulos, A., Harper, C.B., Mavlyutov, T.A., Niranjan, D., Glass, N.R., Cooper-White, J.J., Sibarita, J.B., Choquet, D., et al. (2015). Control of autophagosome axonal retrograde flux by presynaptic activity unveiled using botulinum neurotoxin type a. The Journal of neuroscience: the official journal of the Society for Neuroscience 35, 6179–6194.

Wang, Z., Miao, G., Xue, X., Guo, X., Yuan, C., Wang, Z., Zhang, G., Chen, Y., Feng, D., Hu, J., et al. (2016). The Vici Syndrome Protein EPG5 Is a Rab7 Effector that Determines the Fusion Specificity of Autophagosomes with Late Endosomes/Lysosomes. Mol Cell 63, 781–795.

Weidberg, H., Shvets, E., Shpilka, T., Shimron, F., Shinder, V., and Elazar, Z. (2010). LC3 and GATE-16/GABARAP subfamilies are both essential yet act differently in autophagosome biogenesis. EMBO J 29, 1792–1802.

White, J.G., Southgate, E., Thomson, J.N., and Brenner, S. (1986). The structure of the nervous system of the nematode Caenorhanditis elegans. Philosophical Transactions of the Royal Society of London 314, 1–340.

Wong, E., and Cuervo, A.M. (2010). Autophagy gone awry in neurodegenerative diseases. Nature neuroscience 13, 805–811.

Wu, F., Li, Y., Wang, F., Noda, N.N., and Zhang, H. (2012). Differential function of the two Atg4 homologues in the aggrephagy pathway in Caenorhabditis elegans. The Journal of biological chemistry 287, 29457–29467.

Yin, Z., Pascual, C., and Klionsky, D.J. (2016). Autophagy: machinery and regulation. Microb Cell 3, 588–596.

Yu, Z.Q., Ni, T., Hong, B., Wang, H.Y., Jiang, F.J., Zou, S., Chen, Y., Zheng, X.L., Klionsky, D.J., Liang, Y., et al. (2012). Dual roles of Atg8-PE deconjugation by Atg4 in autophagy. Autophagy 8, 883–892.

Zhang, H., and Baehrecke, E.H. (2015). Eaten alive: novel insights into autophagy from multicellular model systems. Trends in cell biology 25, 376–387.

Zhang, H., Chang, J.T., Guo, B., Hansen, M., Jia, K., Kovacs, A.L., Kumsta, C., Lapierre, L.R., Legouis, R., Lin, L., et al. (2015). Guidelines for monitoring autophagy in Caenorhabditis elegans. Autophagy 11, 9–27.

Zhang, L., Li, J., Ouyang, L., Liu, B., and Cheng, Y. (2016). Unraveling the roles of Atg4 proteases from autophagy modulation to targeted cancer therapy. Cancer Lett 373, 19–26.

Zhao, H., Zhao, Y.G., Wang, X., Xu, L., Miao, L., Feng, D., Chen, Q., Kovacs, A.L., Fan, D., and Zhang, H. (2013). Mice deficient in Epg5 exhibit selective neuronal vulnerability to degeneration. The Journal of cell biology 200, 731–741.

## REFERENCES

Biron, D., Shibuya, M., Gabel, C., Wasserman, S.M., Clark, D.A., Brown, A., Sengupta, P., and Samuel, A.D. (2006). A diacylglycerol kinase modulates long-term thermotactic behavioral plasticity in C. elegans. Nature neuroscience 9, 1499–1505.

Byerly, L., Cassada, R.C., and Russell, R.L. (1976). The life cycle of the nematode Caenorhabditis elegans. I. Wild-type growth and reproduction. Developmental biology 51, 23–33.

Clark, D.A., Biron, D., Sengupta, P., and Samuel, A.D. (2006). The AFD sensory neurons encode multiple functions underlying thermotactic behavior in Caenorhabditis elegans. The Journal of neuroscience: the official journal of the Society for Neuroscience 26, 7444–7451.

Hawk, J.D., Calvo, A.C., Liu, P., Almoril-Porras, A., Aljobeh, A., Torruella-Suarez, M.L., Ren, I., Cook, N., Greenwood, J., Luo, L., et al. (2018). Integration of Plasticity Mechanisms within a Single Sensory Neuron of C. elegans Actuates a Memory. Neuron 97, 356–367 e354.

Hosono, R., Mitsui, Y., Sato, Y., Aizawa, S., and Miwa, J. (1982). Life span of the wild and mutant nematode Caenorhabditis elegans. Effects of sex, sterilization, and temperature. Exp Gerontol 17, 163–172.

