## Supplementary figures and images for "Retrograde Transport and ATG-4.2-Mediated Maturation Cooperate to Remove Autophagosomes from the Synapse"

### Supplementary Materials

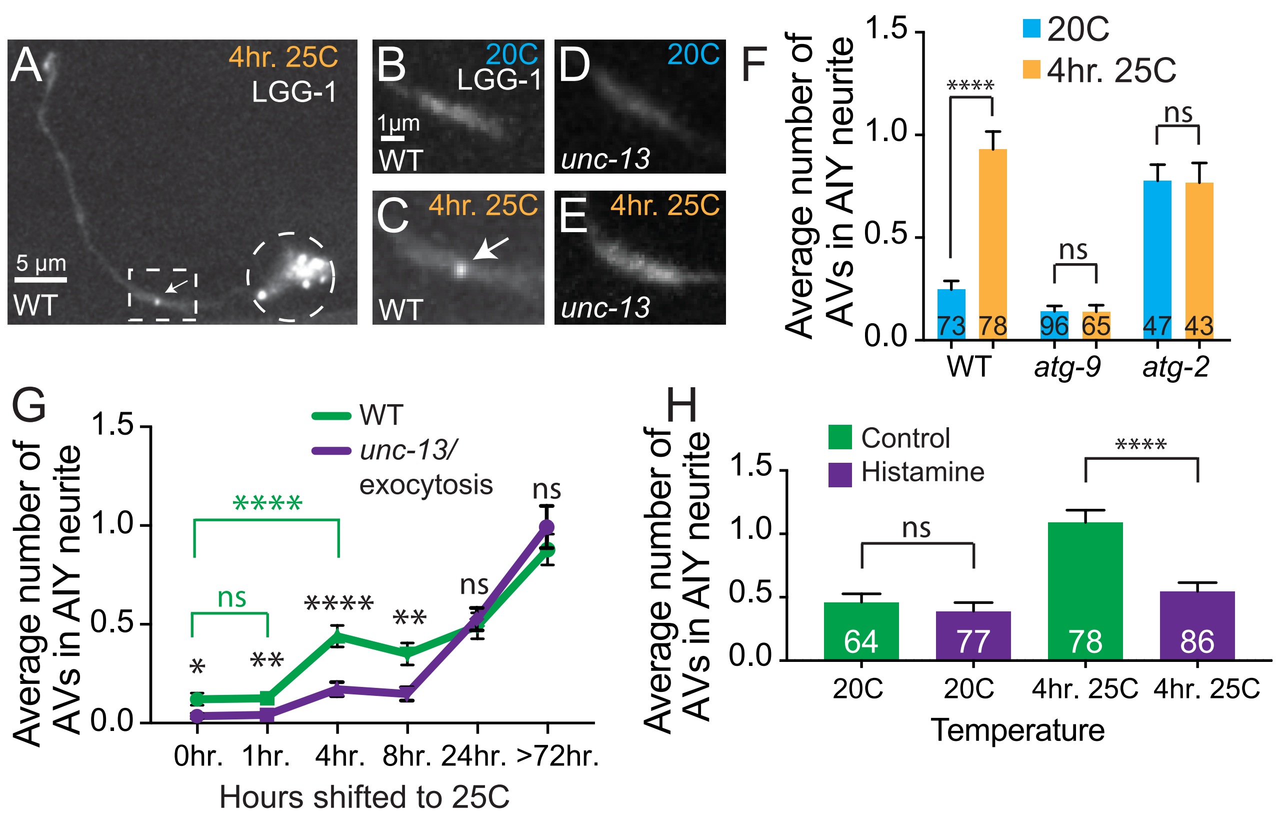

### Supplementary Materials

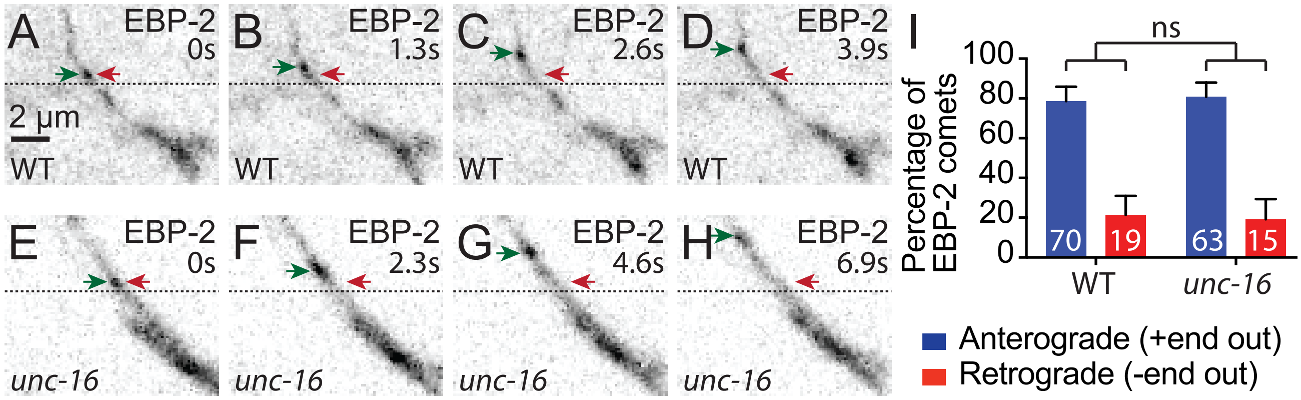

### Supplementary Materials

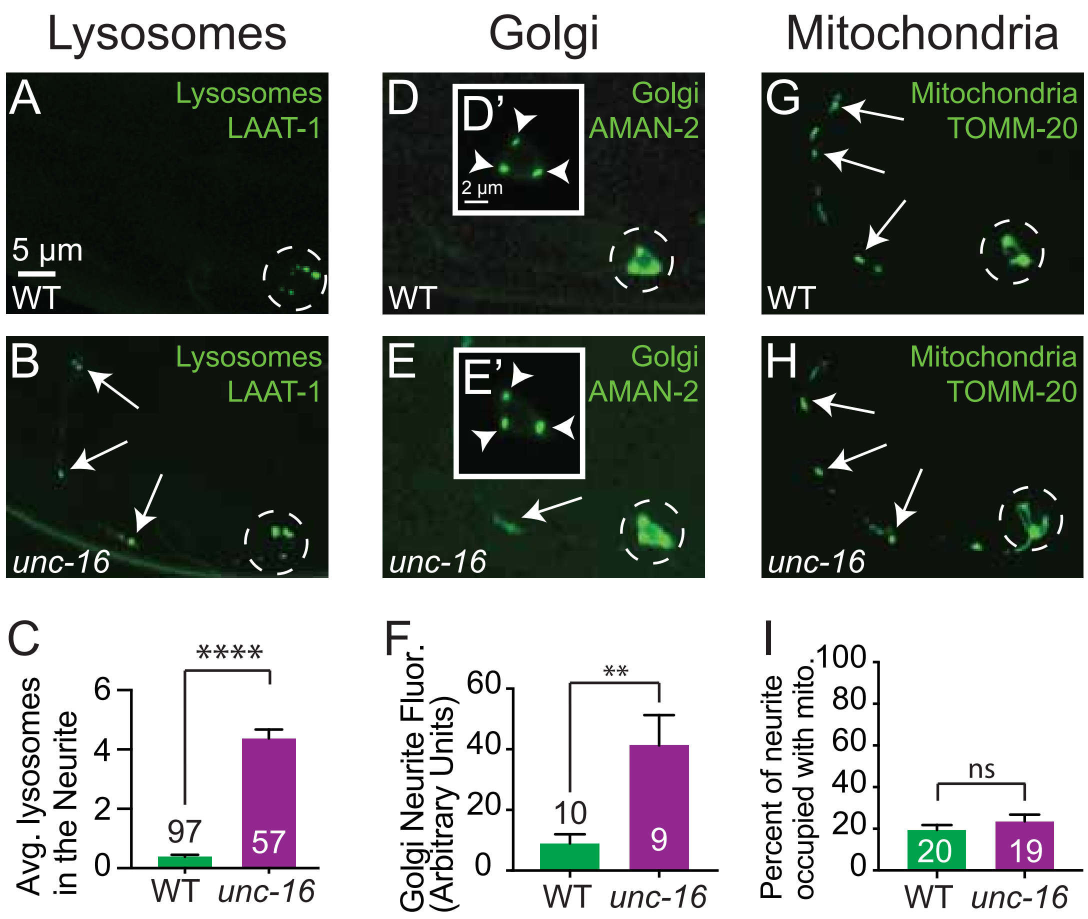

### Supplementary Materials

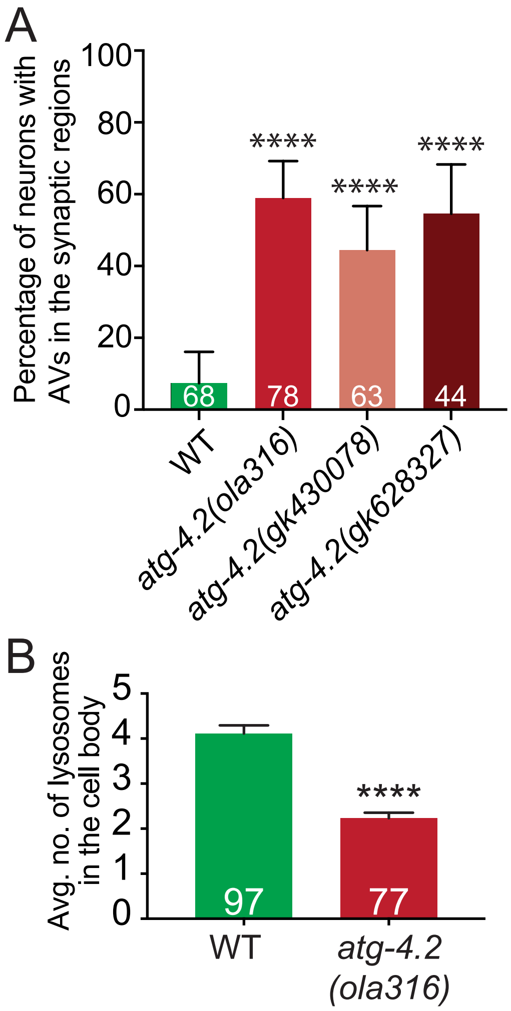

### Supplementary Materials

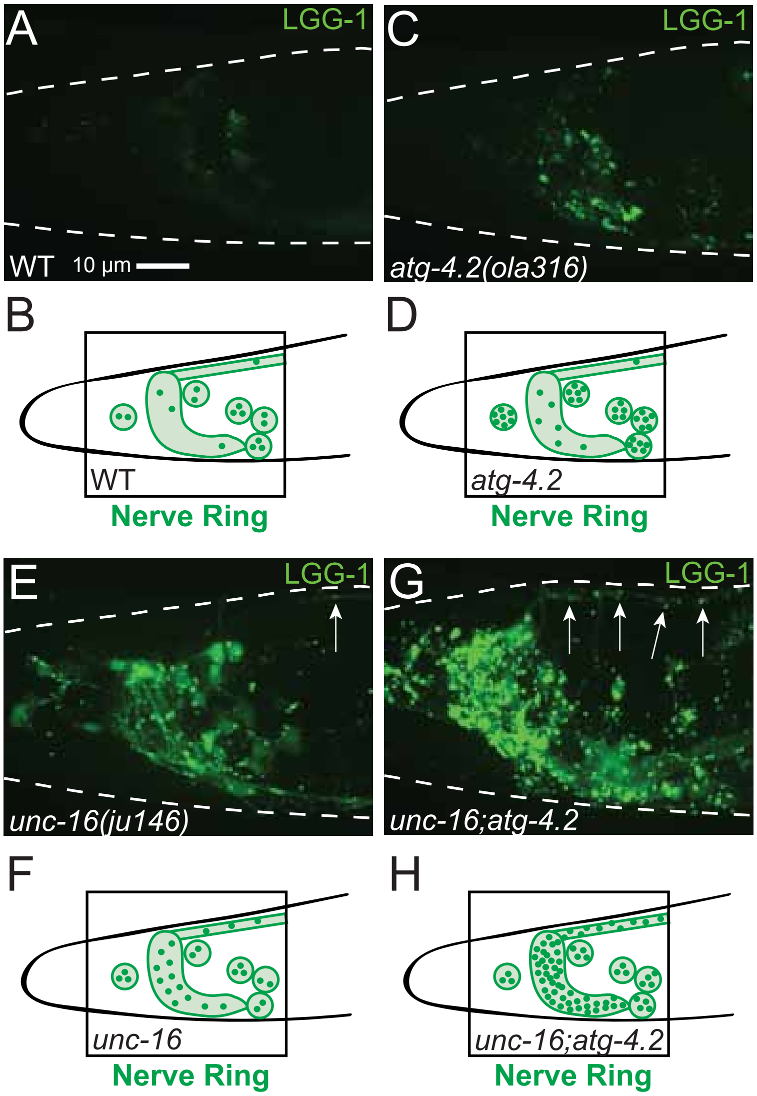

### Supplementary Materials

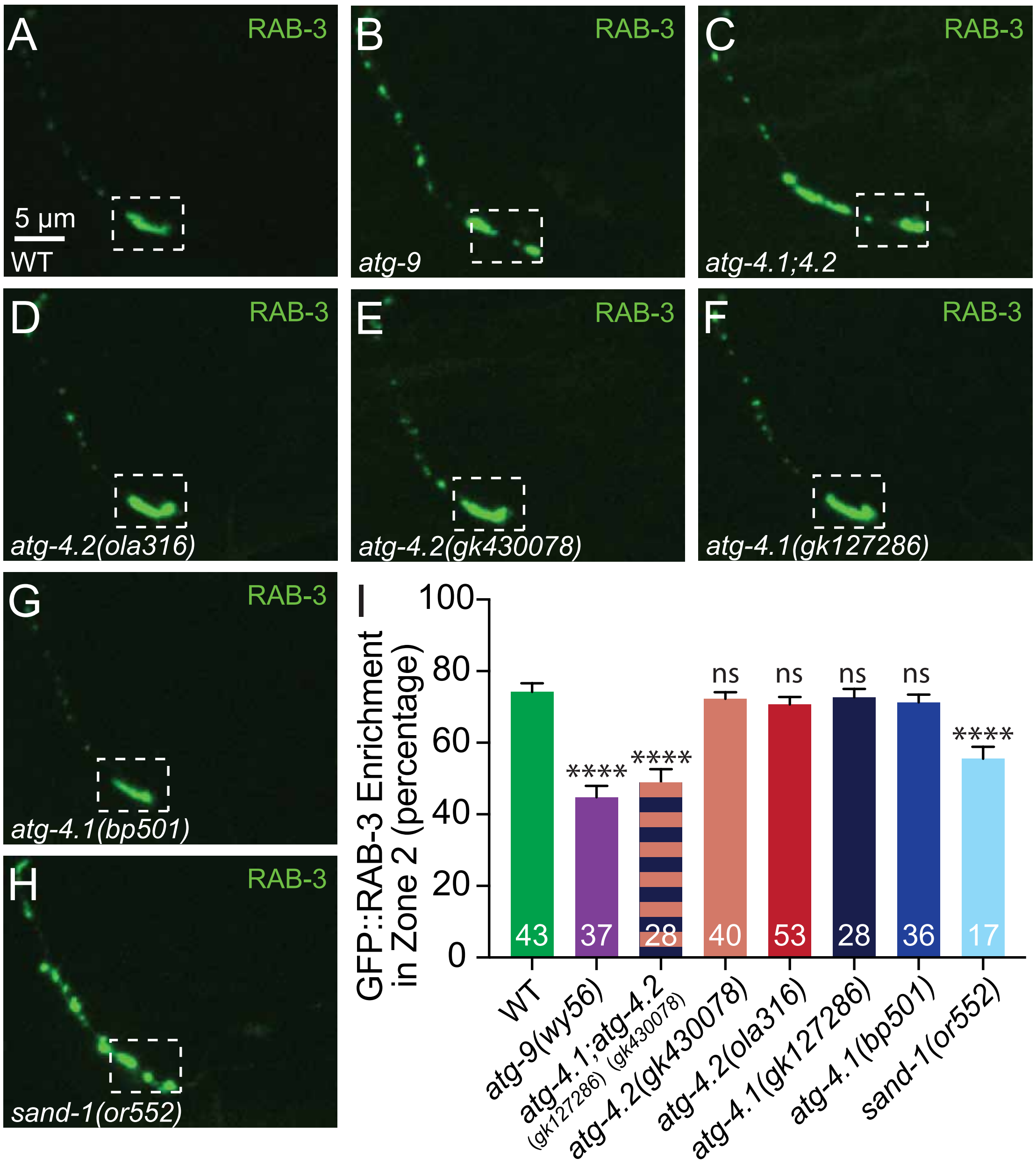
